# Dynamics of Tpm1.8 domains on actin filaments with single molecule resolution

**DOI:** 10.1101/2020.06.15.152033

**Authors:** Ilina Bareja, Hugo Wioland, Miro Janco, Philip R. Nicovich, Antoine Jégou, Guillaume Romet-Lemonne, James Walsh, Till Böcking

## Abstract

Tropomyosins regulate dynamics and functions of the actin cytoskeleton by forming long chains along the two strands of actin filaments that act as gatekeepers for the binding of other actin-binding proteins. The fundamental molecular interactions underlying the binding of tropomyosin to actin are still poorly understood. Using microfluidics and fluorescence microscopy, we observed the binding of fluorescently labelled tropomyosin isoform Tpm1.8 to unlabelled actin filaments in real time. This approach in conjunction with mathematical modeling enabled us to quantify the nucleation, assembly and disassembly kinetics of Tpm1.8 on single filaments and at the single molecule level. Our analysis suggests that Tpm1.8 decorates the two strands of the actin filament independently. Nucleation of a growing tropomyosin domain proceeds with high probability as soon as the first Tpm1.8 molecule is stabilised by the addition of a second molecule, ultimately leading to full decoration of the actin filament. In addition, Tpm1.8 domains are asymmetrical, with enhanced dynamics at the edge oriented towards the barbed end of the actin filament. The complete description of Tpm1.8 kinetics on actin filaments presented here provides molecular insight into actin-tropomyosin filament formation and the role of tropomyosins in regulating actin filament dynamics.

## INTRODUCTION

Actin is a highly conserved and ubiquitous protein found in all eukaryotic cells. With the help of a myriad of actin-binding proteins (ABPs), actin filaments form extensive, highly dynamic, networks which are associated with various structures and functions, including cell division, migration, intracellular transport, cell-cell and cell-matrix adhesion (Pollard 2016). Among the ABPs, tropomyosin is well-known for its roles in the regulation and stabilization of actin filaments. There are multiple isoforms of tropomyosin (~40 in mammals), which are associated with functionally distinct populations of actin filaments (Gunning et al. 2015). Tropomyosin is thought to facilitate the functional specialization by controlling the recruitment of specific sets of actin-binding proteins in an isoform-specific manner (Gunning et al. 2015; Tojkander et al. 2011; Johnson, East, and Mulvihill 2014). Recent studies indicate that the majority of actin filaments in the cell are present as copolymers with tropomyosin (Meiring et al. 2018). Thus, in order to properly understand the variable functions and dynamics of actin filaments, it becomes crucial to elucidate the fundamental molecular interactions between actin and tropomyosin.

Each tropomyosin molecule is a parallel dimeric alpha-helical coiled-coil which covers six or seven actin monomers, depending on whether the isoform is low or high molecular weight, respectively (Khaitlina 2015). Individual tropomyosin molecules bind very weakly to actin (K_a_ ~3×10^3^ M^−1^) (Wegner 1980; Weigt, Wegner, and Koch 1991; Tobacman 2008); however, tropomyosin is able to polymerise along actin, which drastically strengthens its binding affinity through avidity (Wegner 1980; Singh and Hitchcock-DeGregori 2009; Wegner 1979). Tropomyosin molecules bind to the two strands of the double helical actin filament, where they interact in a head-to-tail fashion to form two continuous chains that wrap around the actin filament (Li et al. 2011)(Perry 2001; Khaitlina 2015) as shown schematically in Figure 1B.

**Figure 1.**
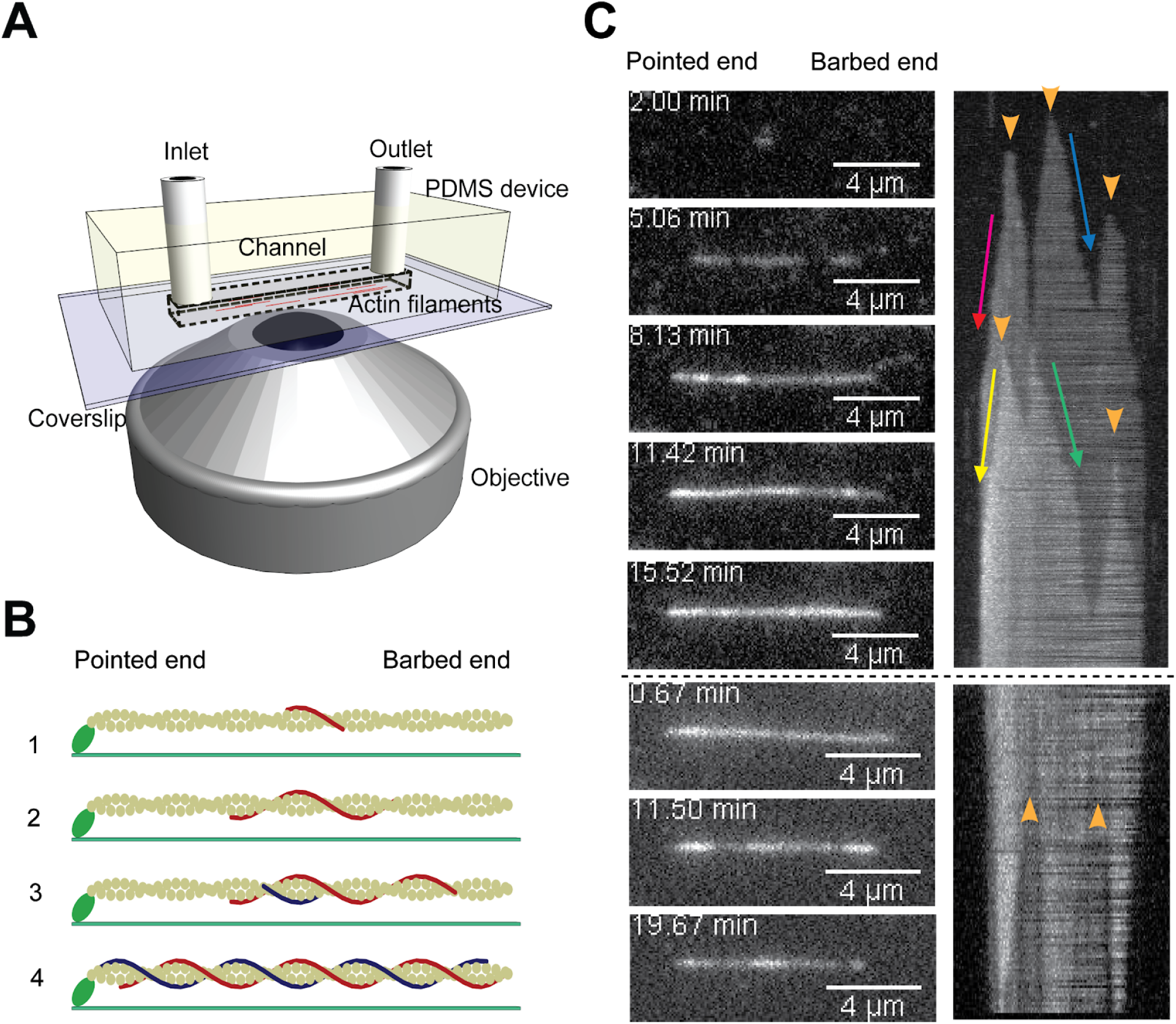
Real time observation of Tpm1.8 assembly and disassembly on actin filaments by TIRF microscopy. **(A)** Microfluidics and TIRF setup. **(B)** Schematic of the steps leading to decoration of an actin filament with tropomyosin: (1) A tropomyosin molecule (red) binds to a random site on one of the two strands on a naked actin filament attached to the surface via spectrin-actin seeds (green). (2) Domain elongation: tropomyosin molecules then bind at adjacent sites and form head-to-tail overlap complexes with the already bound tropomyosin molecule to extend the domain toward both the pointed and barbed ends of the actin filament. (3) Domain appearance and elongation occurs on both strands of the actin filament, as shown by a second (blue) tropomyosin strand. (4) Finally, the double helical actin filament is coated by two tropomyosin chains. The process of dissociation occurs by reversing the process, whereby tropomyosin can dissociate from either of the two ends of the tropomyosin strands located on both strands. **(C)** Kymographs and snapshots from a time-lapse series of a single actin filament showing Tpm1.8 association (top) and dissociation (bottom) after injection and wash-out of mNeonGreen-Tpm1.8, respectively. The two levels of fluorescence intensity correspond to Tpm1.8 domains on either one or both actin strands. Orange arrow heads: nucleation points during association and points where dissociation starts after mNeonGreen-Tpm1.8 wash-out. Arrows indicate the slopes used to measure the elongation rate toward either end.

Cell biological and biochemical techniques have been used extensively to identify different tropomyosin isoforms, their localisation, and the corresponding ABPs they regulate (Bryce et al. 2003; Pathan-Chhatbar et al. 2017; Brayford et al. 2016; Creed et al. 2011; Gateva et al. 2017; Tojkander et al. 2011). Ensemble measurements (solution assays) have shown differences in affinity and cooperativity of tropomyosin isoforms binding to actin and in their effect on actin assembly kinetics (Janco et al. 2016). However, the self-assembly of tropomyosin on actin filaments is a highly stochastic and non-linear process, which is difficult to resolve using ensemble measurements. Basic nucleation and growth models have been proposed (Vilfan 2001) which are able to recapitulate observations from ensemble measurements (Weigt, Wegner, and Koch 1991; Wegner 1980, 1979; Wegner and Ruhnau 1988; Keiser and Wegner 1985; Wegner and Walsh 1981). Parameterisation from these models suggest that binding of tropomyosin to actin filaments involves a slow, initial nucleation step followed by rapid elongation.

Recent developments in reconstituting actin filaments near surfaces for observation by time-lapse fluorescence microscopy have enabled the study of various processes regulating actin dynamics at a molecular level (Jégou and Romet-Lemonne 2016; Shekhar and Carlier 2016; Carlier, Romet-Lemonne, and Jégou 2014) including the interplay between different tropomyosins and other ABPs (Schmidt, Lehman, and Moore 2015; Hsiao et al. 2015; Sckolnick et al. 2016; Christensen et al. 2017; Jansen and Goode 2019). These studies demonstrate the power of this approach for dissecting individual steps of tropomyosin domain nucleation, elongation and shrinkage on actin filaments, which will enable testing and refining our models of these processes. It remains largely unknown to what extent tropomyosin assembly differs between species and between isoforms. A dissection of the common and isoform-specific mechanisms governing the interplay of tropomyosin with actin and their relationship to function, especially in the context of the numerous cytosolic isoforms that are spatially and temporally regulated in mammalian cells (Gunning et al. 2005), will therefore require detailed studies of individual isoforms that are involved in different actin-mediated processes.

The human cytosolic low molecular weight isoform Tpm1.8 is associated with stress fibres and lamellipodia, where it is involved in regulating the highly dynamic process of cell migration (Bryce et al. 2003; Brayford et al. 2016). It exhibits one of the strongest affinities for actin (Moraczewska, Nicholson-Flynn, and Hitchcock-DeGregori 1999), but its assembly and turn-over on actin and how these properties relate to the dynamics of Tpm1.8-containing cellular actin structures remain unresolved. In this study, we used microfluidics and total internal reflection fluorescence (TIRF) microscopy to measure the dynamic interactions of Tpm1.8 molecules with preformed actin filaments *in vitro* at the single filament and the single molecule level. Our data provide a complete experimentally parameterised model of Tpm1.8 assembly and disassembly kinetics on actin, providing molecular insight into the interplay between the two polymer systems.

## RESULTS

### Microfluidics experiments allow for the direct observation of Tpm1.8 dynamics on individual actin filaments in real time

We used the combination of microfluidics and TIRF microscopy to characterise the fundamental molecular interactions underlying tropomyosin binding to actin filaments, as shown in Figure 1A. We immobilized spectrin-actin seeds on the surface of a coverslip that formed the bottom of a microfluidic flow channel. The spectrin-actin seeds anchored actin filaments at their pointed end and allowed for the growth of actin filaments at the barbed end. Each field of view typically contained 50–60 actin filaments (Supplementary Figure 1). After growing actin filaments and ageing them in order to get ADP-F-actin, fluorescent Tpm1.8 was flown into the channel and its binding to the actin filaments was directly imaged using TIRF microscopy. Throughout the acquisition, the filaments were kept aligned parallel to the surface by a constant flow of the solution. This approach allowed the proteins to bind freely to the filaments without any hindrance from the surface. After complete decoration of the filaments, the flow channel was washed with buffer and the dissociation of Tpm1.8 from the filaments was observed. We used recombinant Tpm1.8 fused at its N-terminus to an alanine-serine extension mimicking acetylation (Monteiro et al. 1994) and to mNeonGreen.

Figure 1B summarises the main steps of the actin-tropomyosin interaction that were characterized in this study. The initial assembly intermediates of new tropomyosin domains were detected as the appearance of diffraction limited dots of fluorescent Tpm1.8 molecules. In the range of the concentrations used, multiple such events were detected at different times and locations over the filaments, whereby the number of observable stable domains per filament increased with concentration (Supplementary Figure 2). These dots then elongated in both directions until actin filaments (unlabelled) were fully decorated. The dissociation process, similarly, initiated at multiple points and at different times and the signal disappeared in both directions from the initiation points, which may in large part arise from gaps that remain between adjacent Tpm1.8 domains (Supplementary Figure 3). We were able to resolve the dynamics of Tpm1.8 domains on both strands of actin filaments, as seen by the two levels of fluorescence intensity in Figure 1C. Since Tpm1.8 binds tightly, complete dissociation was not observed in the time scales used for imaging, as seen from the kymograph. Combined, this system enabled us to directly observe the three basic steps of tropomyosin kinetics on both strands of actin filaments: the appearance of Tpm1.8 domains (nucleation), elongation as a result of binding of Tpm1.8 molecules to the edges of domains and shrinkage as a result of dissociation of Tpm1.8 molecules from the edges of domains.

### Tpm1.8 domains have asymmetric dynamics

The kymographs resulting from the microfluidic TIRF binding experiments were used to resolve the Tpm1.8 elongation and shrinkage kinetics towards both ends of the actin filament at a range of Tpm1.8 concentrations (Figure 2). The elongation rates were obtained from the kymographs by taking slopes that correspond to the increase in fluorescence intensity with time as illustrated in Figure 1C. As expected the elongation rates increased linearly with concentration (Figure 2A). Surprisingly, the elongation towards the barbed end (1.19 m.M^−1^.s^−1^) was faster than towards the pointed end (0.64 m.M^−1^.s^−1^) of the actin filament with a ratio of 1.85 between these two rate constants. Similar elongation rates reflecting this asymmetry were measured when we reversed the polarity of actin filaments in the fluid flow by using gelsolin to anchor filaments to the surface from their barbed end (Supplementary Table 1). We also compared the mNeonGreen-Tpm1.8 elongation kinetics on filaments grown from muscle actin (used throughout this work) and on filaments grown from cytoskeletal actin (which is the binding partner of Tpm1.8 in the cell) and found that the domain elongation rates towards both ends of the actin filament were essentially the same (Supplementary Figure 5). These observations showed that the asymmetry in elongation rates was independent of fluid flow or choice of actin isoform.

**Figure 2.**
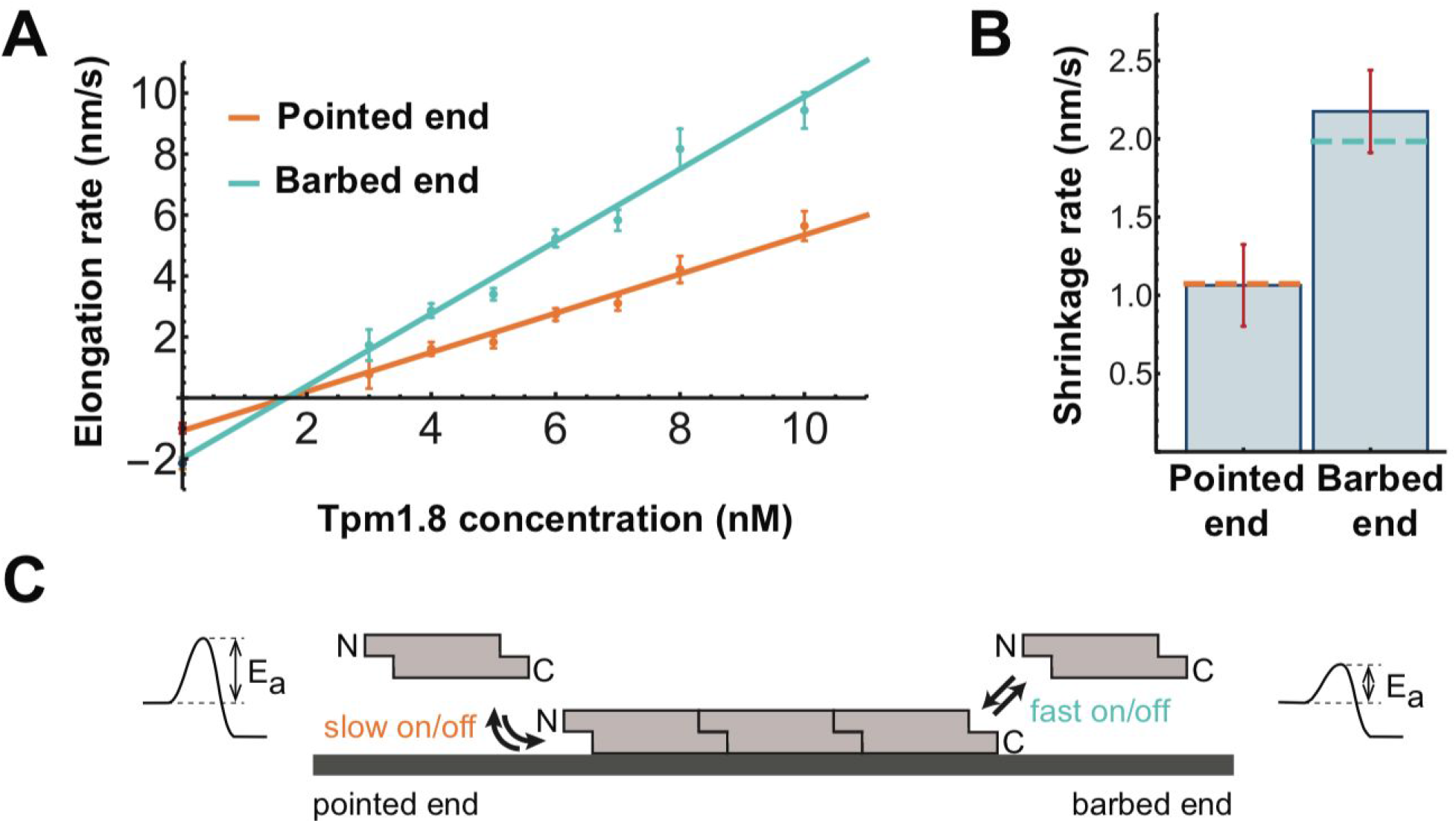
Tpm 1.8 domains grow and shrink faster at the domain edge directed towards the barbed end than the pointed end of the actin filament. **(A)** Tpm1.8 concentration dependence of domain elongation rates toward the barbed and pointed ends of the actin filament. The solid lines represent linear fits of the data, whereby the y-axis intercept of each fit line gives the shrinkage rate for the respective end, the x-axis intercept gives the critical concentration and the slope gives the elongation rate constant. Points represent the mean and error bars represent the standard deviation; *N* (number of filaments [slopes measured towards pointed/barbed end]) = 11 [2/13] (3 nM); 20 [35/66] (4 nM); 40 [73/116] (5 nM); 48 [109/121] (6 nM); 52 [99/110] (7 nM); 38 [73/90] (8 nM); 58 [148/178] (10 nM). **(B)** Comparison of shrinkage rates of Tpm1.8 at the two edges of a Tpm1.8 domain. The values for the shrinkage rates obtained from y-axis intercepts in (A) are represented by dotted lines. *N* (number of slopes) = 78 (pointed end), 111 (barbed end); *p* = 6.6E-14, unpaired Student’s t-test, after Welch’s correction. **(C)** Schematic of the binding of tropomyosin molecules to either end of an existing domain on an actin filament with corresponding potential energy diagrams of the reaction. N, N-terminal end; C, C-terminal end; E_a_, activation energy.

Fluorescence labelling of tropomyosins can impair their function, whereby the effects of different labelling strategies depend on the isoform and experimental system. Fusion of a fluorescent protein tag to the N-terminus of Cdc8, the sole tropomyosin in fission yeast, does not perturb its assembly on actin filaments *in vitro* or localization in the cell (Brooker, Geeves, and Mulvihill 2016) but leads to severe functional defects, in particular misregulation of actin nucleation and cell division (Wu et al. 2003; Brooker, Geeves, and Mulvihill 2016). These defects can be alleviated by using certain cysteine mutants of Cdc8, and Cdc8 labelled at the engineered cysteine can assemble on actin (Christensen et al. 2017). N-terminal fusions of mammalian tropomyosins localize to the expected actin structures and support isoform-specific functions in cells (Appaduray et al. 2016; Tojkander et al. 2011; Sao et al. 2019)also assemble on actin filaments *in vitro* (Gateva et al. 2017). To test whether the N-terminal fluorescent protein affected the kinetics of Tpm1.8 in our assays, we monitored the assembly of an equimolar mixture of tagged and untagged Tpm1.8, which yielded similar elongation rates as observed above (Supplementary Table 1). The fluorescence intensity of actin filaments decorated with the equimolar mixture was 42% of the value for fully tagged Tpm1.8, i.e. close to value expected for equal incorporation of tagged and untagged Tpm1.8. Furthermore, elongation kinetics of the related isoform Tpm1.1 fused to a fluorescent protein or labelled with an organic fluorophore on an internal cysteine were similar to each other and assembly showed the same general features as for mNeonGreen-Tpm1.8, including the asymmetry in elongation rates (Supplementary Figure 4). While it is likely that N-terminal fusions affect some function of tropomyosins, our observations suggest that the fluorescent protein tag only had a minor effect on elongation kinetics and actin affinity in our *in vitro* assays.

Domain shrinkage rates determined from the slopes of signal decrease in the kymographs after wash-out were independent of the mNeonGreen-Tpm1.8 concentration used to decorate the filaments prior to wash-out, as expected (Supplementary Figure 6). Domain shrinkage was also asymmetric (Figure 2B): Release of Tpm1.8 was 1.85 times faster from the domain edge directed towards the barbed end (1.98.10^−9^ m.s^−1^) than at the pointed end (1.07.10^−9^ m.s^−1^) of the actin filament. Overall our single filament data reveal that the polarity of the actin filament with faster growth at the barbed end is reflected by the asymmetrical kinetics of Tpm1.8 domains with faster growth at the C-terminal edge (Figure 2C) with potential implications for allowing Tpm1.8 domain elongation to keep up with actin filament growth (Supplementary Figure 7).

To determine the effective length of a single Tpm1.8 molecule (i.e. the length added to a domain by addition of a Tpm1.8 molecule), we measured the fluorescence intensity per unit length of actin filaments fully decorated with mNeonGreen-Tpm1.8 (Supplementary Figure 8) and related this value to the fluorescence intensity of a single mNeonGreen-Tpm1.8 (Figure 4E, green curve). Using these values, we then found that each Tpm1.8 molecule covered a length of ~33 nm on an actin filament (Supplementary Figure 8), corresponding to six actin monomers (Holmes et al. 1990; Dominguez and Holmes 2011), as expected for low molecular weight tropomyosin isoforms. The effective length can be used to convert the kinetic rates for association and dissociation determined from length changes in the fluorescence images into units of number of molecules per unit time (see Table 1).

**Table 1:**
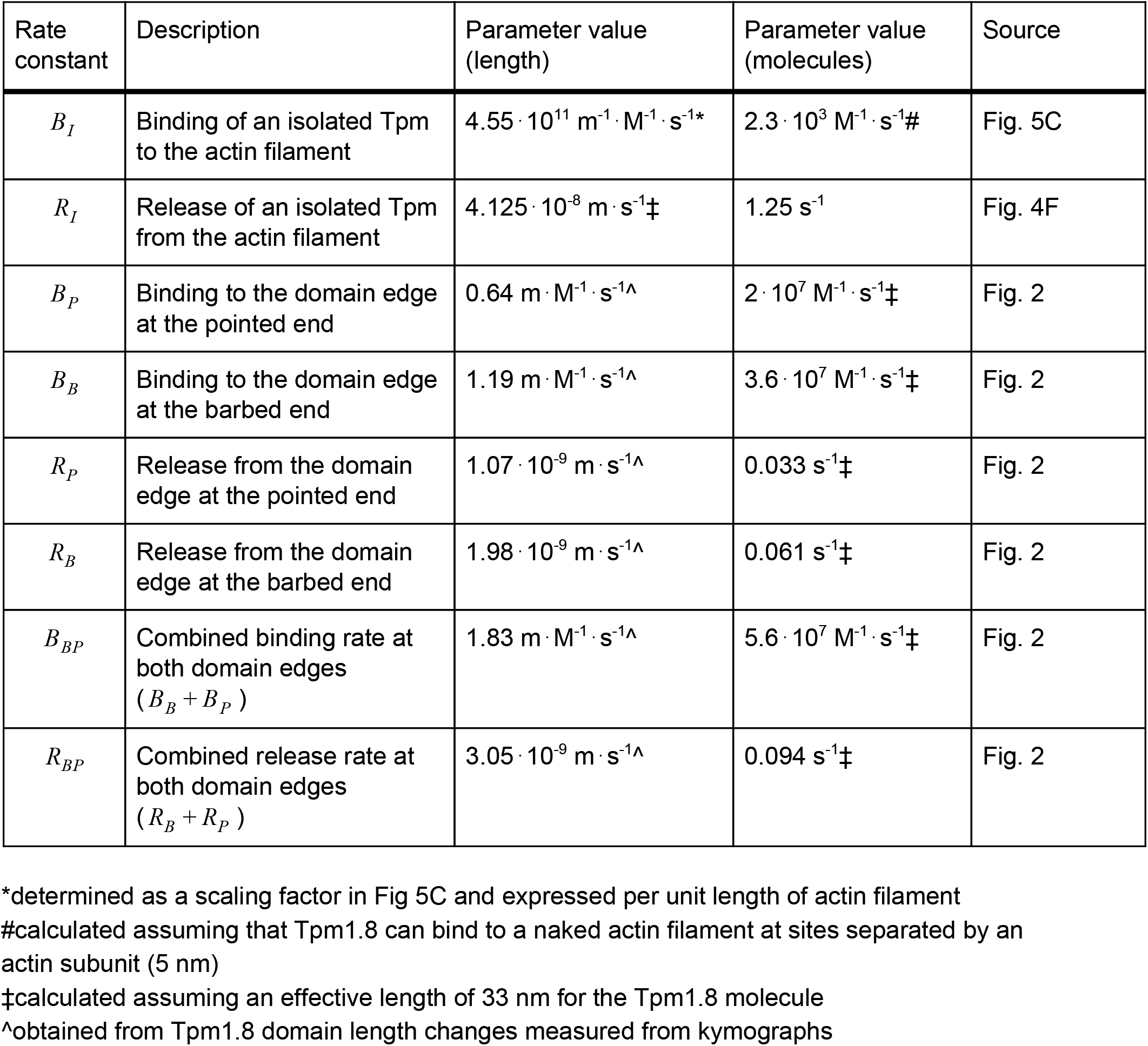
Summary of kinetic rate constants used for modelling.

Finally, we obtained estimates for the critical concentration for the elongation of Tpm1.8 (i.e. the concentration above which binding at domain edges occurs faster than dissociation) from the x-axis intercepts of the fit lines in Figure 2A. As the ratios between the elongation and shrinkage rates are the same for the two domain edges, the critical concentration is also the same towards the two ends of the actin filament (1.65 nM). As a result, there is no concentration where treadmilling (net growth at one domain edge and dissociation at the other) of Tpm1.8 occurs.

### Tpm1.8 decorates the two actin filament strands independently

One unanswered question is whether tropomyosin domains on opposite actin strands can influence each other. We took advantage of our ability to distinguish Tpm1.8 domain kinetics on the two strands to address this question. We measured the following four rates from the corresponding slopes in the kymographs (Figure 1C): (i) first and (ii) second level of increase in intensity at the barbed end of the domain and (blue and green arrow, respectively) (iii) first and (iv) second level of increase in intensity at the pointed end of the domain (red and yellow arrow, respectively). The concentration dependence of these elongation rates (Figure 3A) confirmed the asymmetry towards pointed and barbed ends noted above. However, no significant differences were observed in the elongation rates between the first and second strands in either direction (barbed or pointed). A corresponding analysis of dissociation rates showed that these were the same on stretches of actin filament coated with a Tpm1.8 domain on one or both strands of the actin filament (Figure 3B). Thus, there was no difference in the Tpm1.8 kinetics for the two strands of the actin filament, suggesting that binding of Tpm1.8 to the ends of existing domains occurs independently on each of the two actin strands.

**Figure 3.**
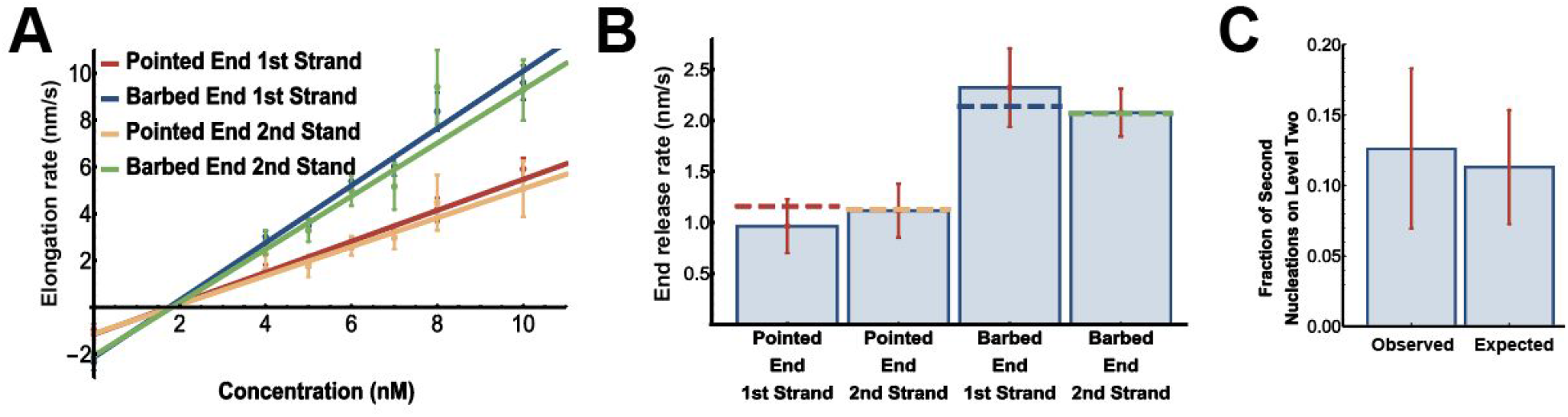
Tpm1.8 kinetics are unaffected by the presence of a Tpm1.8 domain on the opposite strand of the actin filament. **(A)** Plots of tropomyosin domain elongation kinetics toward the barbed and pointed ends of the actin filament as a function of Tpm1.8 concentration, with the data for both strands of the actin filament separated. The lines represent linear fits to the data. Data for pointed end/1st strand, *N* (number of slopes) = 25 (4 nM), 45 (5 nM), 63 (6 nM), 57 (7 nM), 46 (8 nM), 79 (10 nM); pointed end/2nd strand, *N* = 8 (4 nM), 16 (5 nM), 38 (6 nM), 35 (7 nM), 16 (8 nM), 49 (10 nM); barbed end/1st strand, *N* = 38 (4 nM), 71 (5 nM), 73 (6 nM), 72 (7 nM), 58 (8 nM), 89 (10 nM); barbed end/2nd strand, *N* = 18 (4 nM), 23 (5 nM), 32 (6 nM), 23 (7 nM), 15 (8 nM), 59 (10 nM). **(B)** Comparison of shrinkage kinetics of Tpm1.8 toward the two edges of a Tpm1.8 domain, with the data for both strands of the actin filament separated. The values for the dissociation rates obtained from y-axis intercepts in (A) are represented by dotted lines. *N* (number of slopes) = 16 (pointed end, 1st strand), 46 (pointed end, 2nd strand), 20 (barbed end, 1st strand), 82 (barbed end, 2nd strand); *p* = 1.34E-12 (pointed end versus barbed end) and *p* = 0.89 (1st strand versus 2nd strand) using two way ANOVA (Tukey test). **(C)** Frequency of the second Tpm1.8 domain nucleating opposite the already decorated region of the actin filament observed in 198 kymographs (“Observed”). To test for cooperativity of nucleation, this observed frequency of spatial coincidence is compared to the probability of spatial coincidence that is expected when the second domain nucleates at a random location in the undecorated region of the actin filament (“Expected”); one-tailed t-test (*p* = 0.26).

Next, we asked whether the presence of a Tpm1.8 chain on one strand of the actin filament could enhance nucleation of a domain on the opposite strand of the actin filament, as observed for the fission yeast tropomyosin Cdc8 (Christensen et al. 2017). To answer this question, we measured how frequently the second Tpm1.8 domain appeared opposite the first domain versus elsewhere on the actin filament. We also measured the length of the first domain (relative to the length of the actin filament) at the time of second domain appearance and determined the probability that the second nucleation would occur opposite the first Tpm1.8 domain by chance (Figure 3C). We found that nucleations were observed opposite the first Tpm1.8 domain in 25 of 198 kymographs (12.6%), which was not significantly higher than expected by chance (11.3%), suggesting that Tpm1.8 domains have no or minimal effect on second strand nucleation. Overall, we conclude that there is no evidence for the existence of indirect binding cooperativity for Tpm1.8 domain nucleation or elongation between the two strands.

### Tpm1.8 domains grow and shrink one tropomyosin molecule at a time

Using single-molecule imaging, we resolved the kinetics of newly formed Tpm1.8 domains on actin filaments to distinguish whether Tpm1.8 binding and dissociation occurs as monomers or multimeric species. We immobilised actin filaments via multiple (non-specific) anchoring points to the surface (Figure 4Ai). This immobilisation method provided a greater number of filaments per field of view and also facilitated detection of single molecules because the filaments were static (in contrast to filaments grown from a spectrin-actin seed that show movements at the non-tethered end). We then flowed low concentrations of mNeonGreen-Tpm1.8 (3 nM, 4 nM or 5 nM) over the actin to measure initial binding events (appearing as diffraction-limited spots) and early steps of domain elongation (Figure 4Aii), but not full decoration of actin filaments. Fluorescence intensity traces were generated at the sites of initial binding and analysed with a step fitting algorithm that could detect the increase or decrease of the fluorescence intensities as discrete quantal steps (Figure 4C).

**Figure 4.**
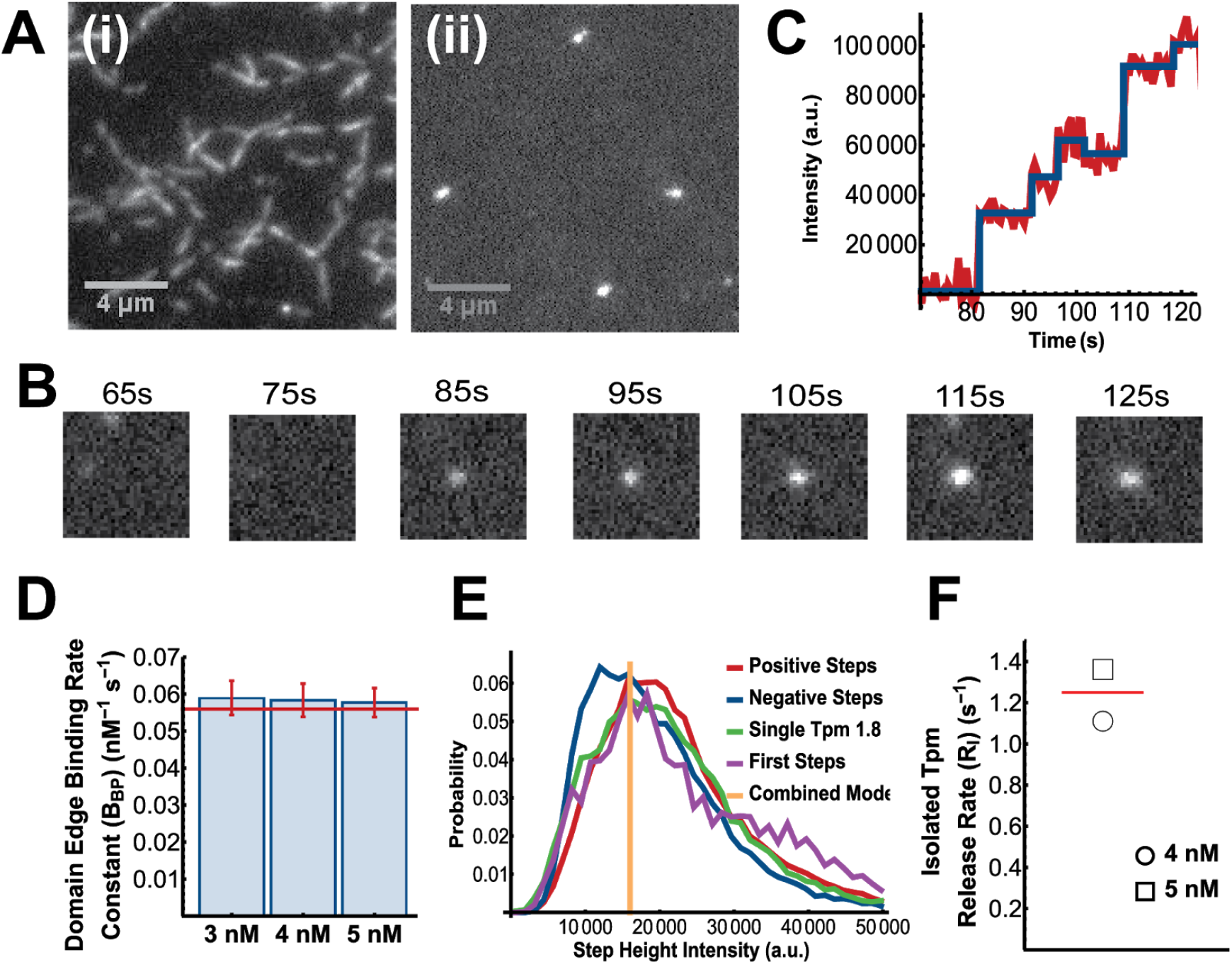
Single-molecule kinetics of Tpm1.8 on naked actin filaments and at the edges of diffraction-limited domains. **(A)** Assembly of mNeonGreen-Tpm1.8 on unlabelled actin filaments attached non-specifically to the surface. (i) TIRF image of actin filaments after complete decoration with mNeonGreen-Tpm1.8 (endpoint of the assembly process). (ii) TIRF image of actin filaments (unlabelled) with short mNeonGreen-Tpm1.8 domains that appeared shortly after injection of mNeonGreen-Tpm1.8 at a concentration of 3 nM. **(B)** Time-lapse images showing nucleation and growth of a diffraction-limited mNeonGreen-Tpm1.8 domain on an unlabelled actin filament. **(C)** Intensity trace of the mNeonGreen-Tpm1.8 segment shown in (B) recorded with an imaging frequency of 2 Hz with step fit (blue line) to identify times of addition/dissociation of Tpm1.8 molecules on the domain undergoing net elongation. **(D)** Combined rate constant for binding at both domain edges (*R*_*BP*_) determined from step fitting of intensity traces of individual segments. The same rate constant determined from kymographs is shown for comparison. *N* (number of positive step times) = 13504 (3 nM), 15002 (4 nM), 14895 (5nM). Errors represent the 95% CI of the residual of the fits **(E)** Distribution of mNeonGreen-Tpm1.8 intensity (green, determined by the intensity of molecules sparsely adhered to a clean glass surface) overlayed with intensity distributions of the first positive step (purple, binding of a mNeonGreen- Tpm1.8 molecule to a stretch of naked actin filament), all positive steps (red; binding events) and negative steps (blue; dissociation events) obtained from the step fitting of intensity traces of individual mNeonGreen-Tpm1.8 domains. *N* (number of events) = 12364 (single mNeonGreen-Tpm1.8 photobleaching), 4767 (first steps), 48168 (positive steps), 44641 (negative steps). **(F)** Estimated release rate (*R*_*I*_) for an isolated mNeonGreen-Tpm1.8 molecule that binds to the actin filament (see Supplementary Figure 6 for details). *N* (number of single molecule events) = 9213 (4nM), 15850 (5nM).

We extracted the mNeonGreen-Tpm1.8 binding kinetics on surface-bound actin filaments by step fitting fluorescence traces recorded during the association phase. The distribution of dwell times between successive positive steps in the fluorescence intensity traces decayed exponentially (Supplementary Figure 9). The association rates obtained by fitting these exponentials for the respective concentrations are shown in Figure 4D and represent the addition of mNeonGreen-Tpm1.8 molecules at both edges of the Tpm1.8 domain. The polymerisation rate constant estimated from this data (5.9.10^7^ M^−1^.s^−1^) was within 10% of the sum of the rate constants for growth at either end determined from the kymographs above (5.6.10^7^ M^−1^.s^−1^). We conclude that the kinetics observed at the filament scale (i.e. linear growth at the ends of domains) is consistent with the molecular scale for newly established domains.

To determine whether Tpm1.8 binds and dissociates from actin filaments solely as a monomer or whether higher order species (dimer, trimers etc.) are involved, we analysed the step heights of the association and dissociation traces. To obtain a baseline for the fluorescence intensity of individual Tpm1.8 molecules, we sparsely immobilized Tpm1.8 on a coverslip and measured individual particle intensities. The intensity distributions for the first positive step (binding of the first Tpm1.8 molecule to a naked region of actin filament), for subsequent positive steps (association) and for negative steps (dissociation) are overlaid with the single molecule intensity distribution of Tpm1.8 molecules in Figure 4E. The strong similarity between all four distributions suggests that Tpm1.8 binds, elongates and dissociates from actin filaments as a monomer.

The observations above confirmed that the first assembly intermediate consists of an isolated (or solitary) Tpm1.8 bound to actin. This complex is thought to be highly unstable but its half-life is unknown. To obtain insight into this process, we analysed distributions of dwell times of fluorescence signals corresponding to single molecules bound to the filament at a given location and obtained a lower limit estimate for the dissociation rate of an isolated mNeonGreen-Tpm1.8 molecule of 1.25 s^−1^ (Figure 4F, see Supplementary Figure 10 for details) .

Taken together our observations suggest that all Tpm1.8 assembly and disassembly steps proceed in units of single molecules, whereby the kinetics of binding and release from the ends of domains occur independent of the length of the domain. Isolated tropomyosin molecules detach from the actin filaments at least an order of magnitude faster than from the edges of domains.

### A model of domain appearance with a stable nucleus containing only two Tpm1.8 molecules is sufficient to recapitulate experimental data

The early stages of tropomyosin domain formation on actin filaments have been difficult to establish. Studies have hinted at an initial, slow process of nucleation followed by a faster process of elongation (Wegner and Ruhnau 1988; Weigt, Wegner, and Koch 1991). However, it is still not known whether this ‘nucleus’ consists of a minimum number of tropomyosins in order to be stable, and what the rate-limiting step of this reaction is. To address these questions, we compare experimental rates of domain appearance with theoretical predictions.

We obtained rates of domain appearance (nucleation events that proceed to form observable, elongating Tpm1.8 domains) by measuring the earliest time at which the first growing mNeonGreen-Tpm1.8 domain was detectable in the kymographs. For this analysis we only considered the appearance of the first domain for each filament, i.e. when the entire filament is available for binding. We assumed that the probability of domain appearance per unit length per unit time is constant, i.e. the time taken for the initial event should be inversely proportional to the length of the filament. Therefore, we multiplied the appearance time by the length of the filament and used this measure to generate survival curves of filaments that remain without Tpm1.8 domain. Exponential fits of these survival curves then provided experimental rates for domain appearance (Supplementary Figure 11), which increased non-linearly with increasing Tpm1.8 concentration (Figure 5C, red data points).

**Figure 5.**
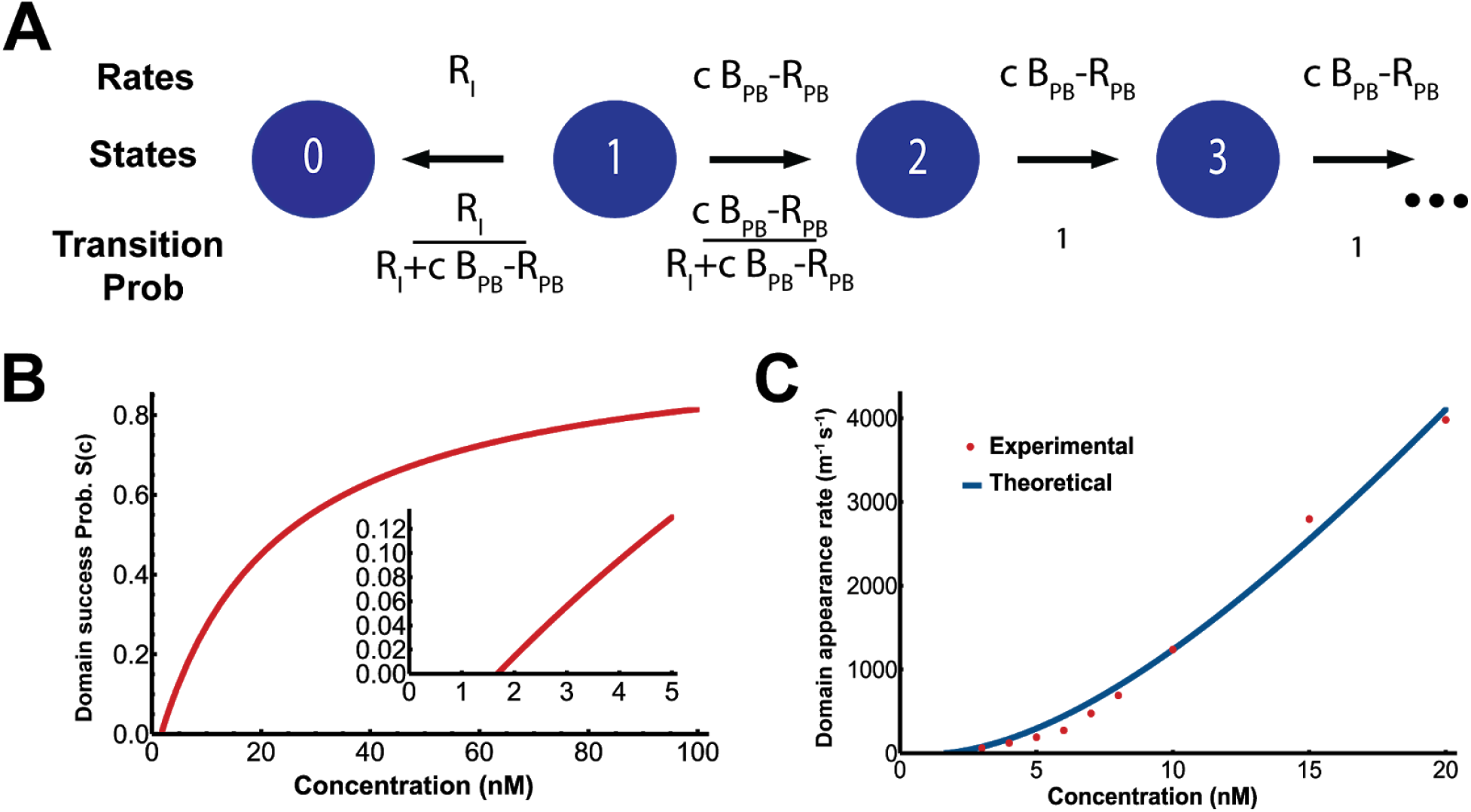
Analysis of domain appearance. **(A)** A simplified model of the growth pathway for a newly formed domain. Single (isolated) Tpm molecules bound to actin are in state *1*. Single molecules that dissociate move to the *0* state at the release rate, *R*_*I*_. Alternatively, a second molecule may bind to the isolated Tpm (*1*), moving it to state *2*. Addition of Tpm to states containing one or more molecules is governed by the elongation rate (*c*.*B*_*BP*_−*R*_*PB*_, where *c* is the Tpm concentration, *B*_*BP*_ and *R*_*PB*_ are the rate constants for binding and release at both edges of the domain). The probability of each reaction is given by dividing each rate by the sum of the rates of reactions out of the relevant state. **(B)** Plot of the probability of domain appearance as a function of concentration calculated using the model in (A). The inset shows the low concentration regime of the curve. **(C)** Domain appearance rate measured experimentally (red dots) from single filaments kymographs (Supplementary Figure 7), and fitted with the nucleation model (blue curve) with the rate constant (B_I_) for binding of an isolated Tpm to the naked actin filament as the only free parameter, yielding a value of B_I_ = 455 nm^−1^ M^−1^ s^−1^.

The kinetic interactions observed in this work are summarized in Figure 6: Reversible weak binding of the first tropomyosin molecule to an undecorated stretch of actin filament is followed by reversible addition of tropomyosin molecules to the edges of the growing domain. The domain appearance rate can be described split into two components: (1) the binding of the first tropomyosin molecule (as a rate *c*.*B*_*I*_) and (2) the probability of continued growth into a tropomyosin strand, i.e. success of nucleation. Together, the domain appearance rate (*k*_*DA*_) can be expressed as:

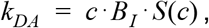

where *c* is the tropomyosin concentration, *B*_*I*_ is the on-rate constant of an isolated tropomyosin molecule binding to the actin filament and *S*(*c*) is the probability of continued growth into an observable tropomyosin domain, i.e. successful nucleation.

**Figure 6.**
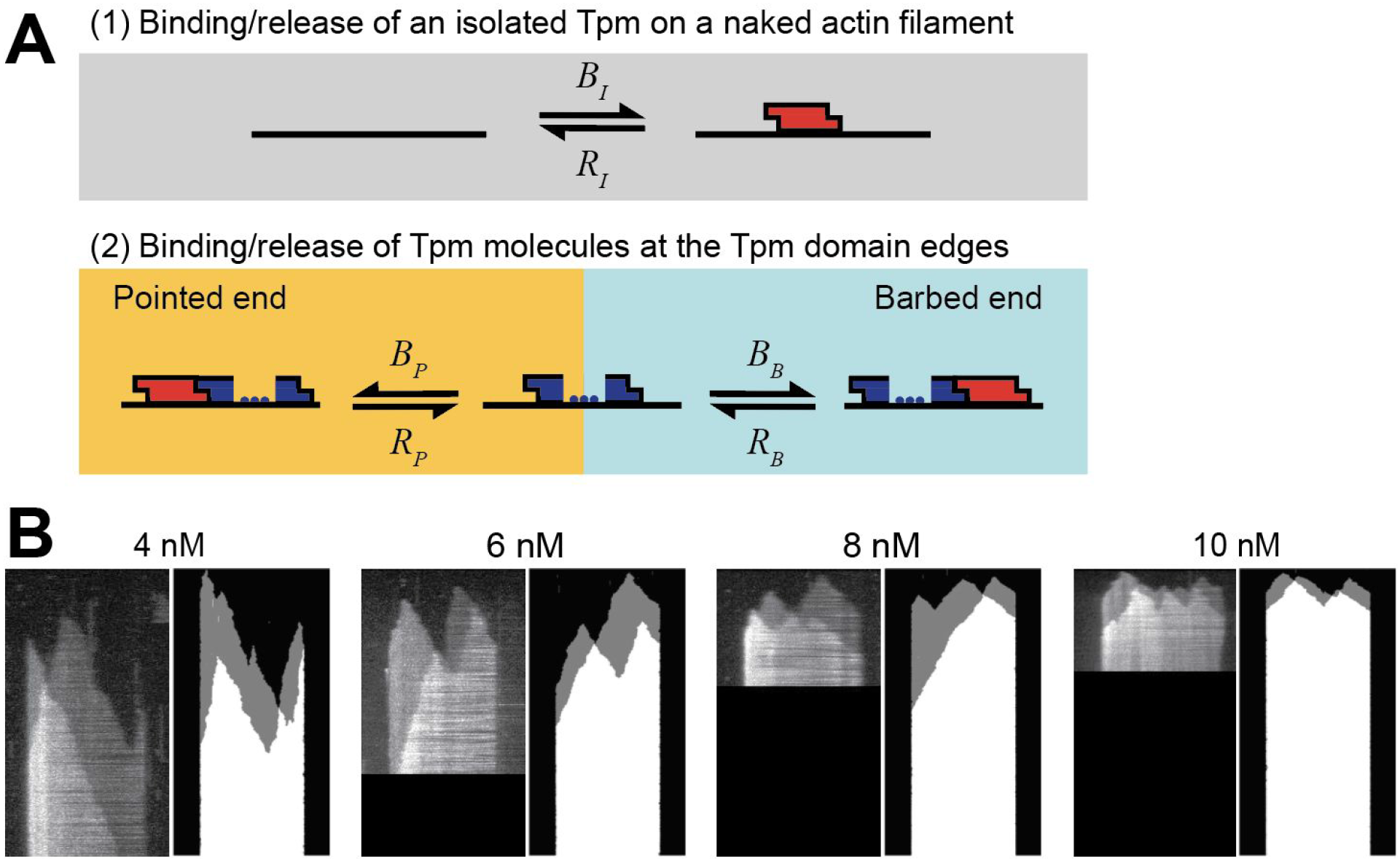
Simulated kymographs of Tpm1.8 assembly on actin filaments obtained with a kinetic model reproduce the features of kymographs from experiments. **(A)** Schematic representation of the model for tropomyosin kinetics on actin filaments. All binding (and dissociation) processes occur by the addition of single Tpm1.8 molecules (shown in red). **(B)** Comparison of experimental and simulated kymographs at a range of Tpm1.8 concentrations. The lengths of filaments from the experiment are 8.449, 8.608, 9.591 and 9.619 μm for 4, 6, 8 and 10 nM Tpm1.8, respectively. The time of acquisition is 55.84, 39.87, 23.34 and 20 min for 4, 6, 8 and 10 nM Tpm1.8, respectively. The simulated kymographs are 8 μm in length. Kymographs are representative of the types of kymographs randomly generated by the simulation.

To explore the interplay between the set of kinetic interactions described in Figure 6 and the probability of continued growth into an observable tropomyosin domain, we converted these interactions into a state space model where each state is denoted by the length of the tropomyosin domain and transitions between states is governed by the effective experimentally measured rates (see Supplementary Information for details). Using this model, we calculated the probability of continued growth (*S*(*c*)) by multiplying all of the transition probabilities from the single tropomyosin state to an infinitely long domain, which gives:

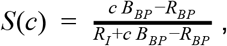

where *c* is the tropomyosin concentration, *B*_*BP*_ and *R*_*BP*_ are the sum of binding and release rate constants, respectively, for the edges of the tropomyosin domain (measured experimentally in Figure 2, See Table 1) and *R*_*I*_ is the rate constant for release of an isolated tropomyosin molecule from the actin filament (measured experimentally in Figure 4, See Table 1). The plot of this function using the experimentally determined parameters (Figure 5B) shows that the probability of an observable domain appearing (i.e. leading to the growth of a Tpm1.8 domain) is zero at tropomyosin concentrations below the critical concentration for tropomyosin domain elongation, and then increases to asymptotically approach 1 at high concentrations, i.e. at high concentrations nucleation is controlled by the binding rate of an isolated tropomyosin molecule (*c*.*B*_*I*_). This calculation shows that in the regime that was probed experimentally (≤20 nM), the nucleation success probability was less than 45%. Figure 5C shows the fit of the nucleation model to the experimental data, whereby *B*_*I*_ is the only free parameter used to minimise the least squares difference with the experimental values providing an estimate of *B*_*I*_ = 455 nm^−1^.M^−1^.s^−1^ (see Supplementary Text and Supplementary Figures 12 and 13 for an alternative method of deriving a value for *B*_*I*_ /*R*_*I*_ from the experimental domain appearance rates).

The domain appearance model is able to reproduce the non-linear increase in the nucleation rate with tropomyosin concentration suggesting that the nucleation process is essentially governed by two sets of reaction rates for (1) binding/dissociation of the first tropomyosin to an undecorated section of an actin filament and (2) binding/dissociation of a tropomyosin molecule at the edges of a domain. We note that in this model the smallest domain at which the latter reaction rates become independent of domain length consists of only two tropomyosin molecules.

### A limited set of kinetic parameters accounts for Tpm1.8 domain dynamics

To test whether the experimentally measured parameters are sufficient to reproduce all features of the observed tropomyosin kinetics, we combined the interactions investigated here (summarised in Figure 6A) into an experimentally parameterized stochastic model (see Table 1 for parameter values and the measurements from which they were derived). In this model we assumed that independently nucleated tropomyosin domains anneal into a single chain when they have sufficiently elongated towards each other to form a contact. In reality, domains presumably only end up annealing into a single chain when they are in register with each other, i.e. the gap between the two domains is multiples of the length of the Tpm1.8 binding site (6 actin subunits) (Supplementary Figure 3). However, given how tightly Tpm1.8 binds, this simplification does not have a significant effect on the kinetics of filament decoration. The kymographs generated using the model at a range of Tpm1.8 concentrations resembled the experimental kymographs (Figure 6B) and faithfully reproduced elongation kinetics and domain appearance rates (Supplementary Figure 14). Overall this comparison confirmed that the steps incorporated into our model were sufficient to predict the experimental kinetics of Tpm1.8 assembly.

## DISCUSSION

Direct observation of Tpm1.8 assembly on actin filaments *in vitro* at the single-molecule and single-filament level with unparalleled resolution enabled us to measure the assembly units and kinetics of Tpm1.8 domain appearance, elongation and shrinkage. On the basis of our observations we propose the following pathway: All association/dissociation steps occur in units of single (dimeric) tropomyosin molecules. A tropomyosin molecule in solution binds to actin with low affinity. Unless additional molecules bind to the first isolated molecule to form a stable domain on actin, it rapidly dissociates into solution. Binding and release occur at both edges of the tropomyosin domain, but more rapidly at the domain edge oriented towards the barbed than the pointed end of the actin filament. Overall domain elongation rates are the same for diffraction-limited (<10 molecules) and microscopic (10 to >100 molecules) domains, i.e. there is no apparent dependence of elongation on domain length. A stochastic model incorporating these reactions is sufficient to simulate experimentally observed tropomyosin kinetics, including rates of domain appearance, suggesting that there are no additional steps governing this process.

The complex between an isolated Tpm1.8 molecule and the actin filament is short-lived (half-life of ~0.6 s or less) and has a low affinity (*K*_*d*_ = *R*_*I*_/*B*_*I*_ ≈ 500 μM), while a tropomyosin molecule bound at the edge of a domain has a longer half-life (11 s and 21 s at the barbed and pointed ends, respectively) and profoundly higher affinity (given by the critical concentration of 1.65 nM). Our analysis suggests that this pronounced increase in stability is largely achieved upon binding of a second tropomyosin to the first (solitary) molecule on actin. That is to say, we propose that the minimal domain (or nucleus) with sufficient stability such that it can grow or shrink with approximately the same kinetics as much longer domains consists of just two molecules. This model is supported by the single-molecule data, where dissociation of domains with two or more tropomyosin molecules was not prominent. Indeed, a domain consisting of two Tpm1.8 molecules almost wraps around the actin filament, and it is tempting to speculate that shorter tropomyosins (such as those found in yeast) would require three molecules to form a stable domain. Nucleation occurs at all tropomyosin concentrations above the critical concentration (1.65 nM) and ultimately leads to the complete decoration of the actin filament (as long as tropomyosin is in excess of actin), consistent with the cooperativity of tropomyosin assembly on actin (Wegner 1979). However, at low concentrations this process is slow because most isolated tropomyosins binding to a naked actin filament fall off before they are stabilised by a second molecule to form growing domains (Figure 4B). Tuning the affinity of an isolated tropomyosin for actin provides a potential control mechanism, whereby increasing the affinity would accelerate domain appearance (nucleation), while a decrease would make nucleation so improbable that it would not occur on relevant timescales.

Tropomyosin cooperativity due to the overlap between N- and C-termini of adjacent molecules in the chain is considered an important characteristic of F-actin network formation, organization, and ABP sorting (Gunning et al. 2015). In addition, it has been proposed that long-range binding cooperativity may arise from the interaction between the two tropomyosin chains binding to the two strands of the actin filament (Schmidt, Lehman, and Moore 2015)(Christensen et al. 2017). Thus, the presence of a tropomyosin chain on one strand of the actin filament may enhance binding of tropomyosin on the opposite strand, which could result in enhanced domain nucleation and/or elongation. We tested for the existence of this additional cooperativity in our data by comparing the elongation and shrinkage rates for the two tropomyosin chains, represented by the two levels of fluorescence intensity in the kymographs. From these measurements, we observed no difference in the kinetics for the two Tpm1.8 chains. We also observed no increase in domain nucleation frequency opposite of an existing Tpm1.8 strand. Taken together our observations indicate that Tpm1.8 binds independently on the two strands of actin filaments (Figure 3).

While the binding affinity of Tpm1.8 is the same at both edges of a domain, we discovered that the on-/off-kinetics are up to 2-fold faster at one edge compared the other, as previously observed (albeit to a lesser extent) for the yeast tropomyosin Cdc8 (Christensen et al. 2017). Tropomyosin chains are oriented on the actin filament such that N-terminus of tropomyosin is directed towards the pointed end while the C-terminus is directed towards the barbed end (Orzechowski et al. 2014) (Figure 2C). The asymmetry in rates at opposing edges could result from an increased activation energy barrier for binding and dissociation at the domain edge with exposed N-terminus compared to the domain edge with exposed C-terminus (Figure 2C). Such an increase in activation energy could arise if the N-terminus of a tropomyosin bound to actin is poorly accessible, e.g. due to reduced conformational plasticity required for forming an overlap junction (Orzechowski et al. 2014). It should be noted that the N-terminal fluorescent protein tag used here can affect the molecular interactions between tropomyosins and between tropomyosin and actin, but the principle characteristics of mNeonGreen-Tpm1.8 kinetics, including asymmetry, were also observed for mixtures of tagged and untagged Tpm1.8 as well as for Tpm1.1 labelled at the cysteine residue at position 190 with an organic fluorophore. The physiological relevance of the asymmetry in kinetics at opposing domain edges, if present for native tropomyosins in the cell, is unclear, but may be relevant for a potential role of Tpm1.8 in regulating actin dynamics (e.g. in lamellipodia (Brayford et al. 2016)) or its interplay with other actin-binding proteins.

Several features of mNeonGreen-Tpm1.8 assembly, such as nucleation and bidirectional growth of multiple domains on both strands of the actin filament leading to full decoration have also been observed for chemically labelled isoforms from a range of species including mammalian Tpm1.1 (produced recombinantly (<u>Nicovich et al. 2016; Janco et al. 2018</u>) or purified with other isoforms from muscle (Schmidt et al. 2015)), fission yeast Cdc8 (Christensen et al. 2017<u>; Palani et al. 2019</u>), or non-muscle *Drosophila* Tm1A (Hsiao et al. 2015), whereby the latter preferentially nucleates domains on ADP-bound regions of the actin filament near the pointed end, which then elongate towards the pointed end. Other characteristics have either not been measured for other isoforms or show different behaviours, such as binding cooperativity for tropomyosin domain nucleation on opposite strands on the actin filament (absent for Tpm1.8 but observed for Cdc8 (Christensen et al. 2017)).

In this work, we have characterised the molecular interactions of Tpm1.8 binding to actin. This provides a platform to unravel how tropomyosin decoration of actin is controlled within human cells. Similar approaches can be applied to other isoforms of tropomyosin to further identify characteristics common to isoforms associated with similar functions. Further, the addition of regulatory proteins or drugs will allow for the identification of the specific molecular interaction being affected.

## MATERIALS AND METHODS

### Constructs

The coding sequence for a fusion protein of mNeonGreen and mRubyII Tpm1.8 was cloned into a pET28a(+) vector using the EcoRI and XhoI sites and additional features were introduced using site-directed mutagenesis. Expression of the final construct yields a protein (referred to as mNeonGreen-Tpm1.8 and mRubyII-Tpm1.8) with N-terminal His_6_-tag followed by mNeonGreen and a peptide linker (GGGSGGGSGTAS) fused to the N-terminus of Tpm1.8. The coding sequence for a fusion protein of mCherry Tpm1.8 was cloned into a pHAT vector using the BglII and HindIII sites. Expression of the final construct yields a protein (referred to as mCherry-Tpm1.8) with N-terminal His_6_-tag followed by mCherry and a peptide linker (SGLRSGGGGSGGGGSGTAS) fused to the N-terminus of Tpm1.8.

### Protein expression and purification

The fusion proteins mNeonGreen-Tpm1.8 and mRubyII-Tpm1.8 were expressed in Rosetta pLysS cells grown in 1 L of LB broth containing 50 μg/mL kanamycin at 37°C. The fusion protein mCherry-Tpm1.8 was expressed in BL21 DE3 star cells grown in 1 L of LB broth containing 0.1 mg/mL ampicillin at 37°C. Protein expression was induced with IPTG (1 mM) when the O.D_600_ reached 0.6 and the cells were grown for an additional 4 h. Cells were pelleted by centrifugation (F10S-6x500Y rotor, Sorvall RC6) at 8000 rpm for 10 min at 4°C. The cell pellet was stored at −40°C until use. The pellet was resuspended in lysis buffer (20 mM Tris-HCl pH 7.8, 500 mM NaCl, 1 mM NaN_3_, 5 mM imidazole, 2 mM MgCl_2_, 0.1% Triton X100, 2% glycerol, 1 mM phenylmethylsulfonyl fluoride (PMSF) and Roche-Hitachi EDTA free protease inhibitor tablet) and sonicated (8 min, 15s on, 15s off). The lysate was centrifuged (R18A rotor, Hitachi VX22N) at 15000 rpm for 60 min at 4°C. The supernatant was filtered using a 0.45 μm syringe filter and loaded onto a 5 mL Hi-Trap column (GE Healthcare Life Sciences) equilibrated with lysis buffer. The column was washed with 50mL buffer containing 30 mM imidazole, 500 mM NaCl, 20 mM Tris, 2 mM MgCl_2,_ 1 mM NaN_3_, pH 7.8 followed by 75mL buffer containing 100 mM imidazole. The protein was eluted using a buffer containing 250 mM imidazole. The fractions with the protein were pooled and further purified by size exclusion chromatography using a 16/60 superdex 75pg column (GE Healthcare Life Sciences) equilibrated with a buffer containing 10 mM Tris pH 7.8, 150 mM NaCl, 1 mM EDTA, 2 mM DTT, 1 mM NaN_3_ and 1% sucrose. The fractions containing protein were pooled, snap frozen in aliquots using liquid nitrogen and stored at −80°C.

### Microfluidics setup

Microfluidic devices with five channels (11 mm × 0.8 mm × 0.06 mm, L×W×H) were prepared from polydimethylsiloxane (PDMS) by replica moulding and ports for tubing (inlet/outlet) were punched at the channel ends using a biopsy punch (diameter 0.7 mm). The device was washed with isopropanol and MilliQ water before use. Coverslips (Marienfeld superior No. 1.5H, 24 mm × 60 mm) were sonicated in filtered 100% ethanol for 30 min, washed with MilliQ water, sonicated in filtered 1 M NaOH for 30 min and washed with MilliQ water. The coverslips were blown dry under a stream of filtered nitrogen and then further dried at 70°C. The PDMS replica and the coverslip were treated with an air plasma for 3 min at 700 torr. The channel device was assembled by pressing the PDMS replica onto the coverslip and annealing the device at 70°C for 4-5h.

### TIRF microscope setup

TIRF data were acquired using a custom-built TIRF microscope based around an ASI-RAMM frame (Applied Scientific Instrumentation) with a Nikon 100× CFI Apochromat TIRF (1.49 NA) oil immersion objective. Lasers were incorporated using the NicoLase system (Nicovich et al. 2017). Images were captured on two Andor iXon 888 EMCCD cameras (Andor Technology Ltd) and 300 mm tube lenses were used to give a field of view of 88.68 μm × 88.68 μm at Nyquist sampling frequency (86 nm per pixel). On our system we typically use a power density of ~1–3 W cm^−2^ (measured at the objective with the laser beam normal to the surface of the coverslip)

### Channel preparation and surface chemistry for single filament experiments

#### Anchoring of filaments at the pointed end using spectrin-actin seeds

Spectrin-actin seeds were prepared as described previously (Casella, Maack, and Lin 1986), diluted into PBS to final spectrin-actin concentration of 30 nM, injected into the channel and allowed to incubate for 4 min. The channel was washed with PBS and incubated with a solution of PLL-PEG (SuSoS AG) in PBS (1 mg/mL) for 30–60 min. The channel was connected to the syringe pump via the outlet tubing and washed with Buffer F (5 mM Tris-HCl pH 7.8, 100 mM KCl, 0.2 mM ATP, 1 mM DABCO, 0.1 mM CaCl_2_, 0.1% NaN_3_, 10 mM DTT, 1 mg/mL BSA, 1 mM MgCl_2_, 0.2 mM EGTA). The channel was then filled with a BSA solution (10 mg/mL in PBS) and incubated for 5 min followed by a wash with buffer F. A solution of G-actin in buffer F (0.8 μM, rabbit skeletal muscle actin, Hypermol EK, Germany) was flowed into the channel and actin filaments were grown from the surface-immobilised spectrin-actin seeds for 6 min. Actin was then flowed through the channel at its critical concentration (0.1 μM) for 15 min, to prevent further polymerisation or depolymerisation and allow for complete phosphate release.

#### Anchoring of filaments at the barbed end using gelsolin

Chambers were prepared as described above with the following modifications. The glass surface was passivated using PLL-PEG-biotin (1 mg/mL, > 1 h). The surface was further passivated with BSA (5 %, 10 min) and casein (Hammarsten Bovine, 5 mg/mL, 10 min). Neutravidin was injected (5 μg/mL, 10 min) followed by biotin-gelsolin (5 nM, 3 min, purified as described in (Wioland et al. 2017)) . Unlabelled F-actin was prepolymerized (8 μM, > 1h), diluted to 0.8 μM and injected in the microfluidic chamber to bind to surface-immobilised gelsolin.

### Single filament imaging of Tpm1.8 association and dissociation

A solution of mNeonGreen-Tpm1.8 in Buffer F (containing 0.1 μM actin to avoid depolymerisation of actin filaments) was flowed through the channel at a flow rate of 30 μL/min and association on actin filaments attached to the glass surface was recorded by time-lapse TIRF imaging (488 nm laser, 20 mW, 30 ms exposure time) with a frame rate depending on the concentration of mNeonGreen-Tpm1.8. Dissociation was initiated by flowing Buffer F (containing 0.1 μM actin) through the flow channel while recording TIRF images with a frame rate of 0.1 Hz.

### Channel preparation and surface chemistry for single molecule experiments

The channel was incubated with a solution of PLL-PEG (1 mg/mL in PBS) for 20 min, connected to the syringe pump via the outlet tubing and washed with buffer F. A solution of 0.8 μM actin in Buffer F was flowed into the channel and incubated for 2 min. During this period actin filaments polymerised in solution and adhered non-specifically to the surface of the coverlip. A wash with Buffer F containing 0.1 μM actin was then given.

### Single molecule imaging of Tpm1.8 association and dissociation

A solution of mNeonGreen-Tpm1.8 (3, 4 or 5 nM) in Buffer F (containing 0.1 μM actin) was flowed through the channel and association on actin filaments attached to the glass surface was recorded by time-lapse TIRF imaging (488 nm laser, 45 mW, 50 ms exposure time). The length and frame rate of the acquisition were adjusted depending on the concentration to obtain a sufficient number of nucleation and elongation steps without significant photobleaching.

### Image analysis at the level of single filaments

#### Elongation and dissociation rates

Kymographs of Tpm1.8 association and dissociation were generated using the FIJI kymograph plugin. The slopes of the kymographs were fitted using linear regression in Wolfram Mathematica to obtain elongation and dissociation rates.

#### Initial domain appearance rate

We assumed that the probability of binding anywhere on the filament is the same such that the time taken for the first nucleation to occur is inversely proportional to the length of the filament. Hence, the product of time of the first nucleation event (*t*) and length of the filament (*l*) is a constant. We plotted survival curves for the number of actin filaments without nucleation event as a function of the *l*×*t* constant, which decreased exponentially. Fitting these exponentials gave the nucleation half-lives for different concentrations of Tpm1.8.

### Nucleation cooperativity analysis

The experimental frequency of a second domain nucleation occurring opposite the first domain was determined from kymographs. For each kymograph we also measured the fraction (*f*) of the filament covered (on one strand) by the first Tpm1.8 domain at the nucleation time of the second domain. The probability that the second nucleation would occur opposite the first domain is then given by *f/(2-f)*. The probability was further corrected for a slight bias in detecting nucleation sites closer to the anchor site (possibly arising from actin filament dynamics and/or anchoring artefacts) in our data. Without this bias correction, the probability of spatial coincidence is 0.106±0.02 (mean±s.d.) for nucleation at random locations, which is not significantly lower than the experimental frequency (*p* = 0.16).

### Image analysis at the single molecule level

Image stacks were analysed using the JIM Immobilized Microscopy Suite to extract intensity traces at the sites corresponding to nucleation events (https://github.com/lilbutsa/JIM-Immobilized-Microscopy-Suite). Intensity traces were divided by the intensity of a dimeric mNeonGreen-Tpm1.8 molecule (equal to twice the intensity measured for mNeonGreen). Stepfitting of traces was achieved using the findchangepts function in Matlab with a threshold of 0.03. Histograms of the intensity step heights for nucleation (first step), elongation (subsequent positive steps) and dissociation (negative steps) of Tpm1.8 were generated using Wolfram Mathematica. Cumulative probability functions of the time before a positive/negative step with fitted with single exponential decays to obtain elongation/dissociation rates, respectively. Dividing the association rates by the concentration gave the association rate constant.

### Stochastic Model

The stochastic model was solved using a custom program written in Mathematica 10.2 (Wolfram). The script used the Gillespie algorithm to generate statistically correct trajectories (Gillespie 1976, 1977). A list containing the ends of each tropomyosin domain was used to calculate the probability of each reaction. The code for this program is freely available at (https://github.com/lilbutsa/Tropomyosin_Stochastic_Model).

## ACKNOWLEDGEMENTS

Funding (T.B.) from the Australian Research Council and National Health and Medical Research Council of Australia. We thank Peter Gunning for reagents and discussions and Richard Morris for discussions. We thank Berengere Guichard for help with protein purification and molecular biology.

## AUTHOR CONTRIBUTIONS

IB: conceptualization, methodology, investigation, formal analysis, writing (original draft); HW: investigation, validation, formal analysis; MJ: methodology, resources; PRN: resources; AJ: methodology; GRL: conceptualization, methodology; JW: conceptualization, software, methodology, formal analysis, writing (original draft); TB: conceptualization, methodology; all authors: writing (review and editing)

## Supplementary text

### Mathematical analysis of tropomyosin domain appearance

The process of growth from an initial isolated Tpm bound to the actin filament can be modelled as shown schematically in Supplementary Figure 9A. In this model, the number of each state denotes a Tpm domain that contains *i* (dimeric) Tpm molecules. As all domains begin as a single molecule, the nucleation process starts in state *1*. From this state, the molecule will either fall off to state *0* at the release rate, *R*_*I*_, or grow by the addition of a second molecule to state 2, at a rate *c B*_*BP*_. We note that in this model, state *2* represents the smallest Tpm1.8 domain at which the on- and off-rates at the ends become independent of domain length, i.e. the nucleus in this model contains two Tpm1.8 molecules. Each higher state can either grow (*i* → *i* + 1) or shrink (*i* → *i* − 1) by binding or releasing Tpm at either edge of the domain, respectively.

Ultimately, we are interested in calculating the probability that a domain successfully elongates rather than getting the time scale on which it occurs. As such, it makes sense to consider the transition probabilities rather than rates. Summing over the probability paths will then give the probability that a domain takes that route. The probability of transitioning from one state to its neighbouring state is found by dividing the corresponding rate by the sum of all rates out of the state.

The probability of a stable domain growing from an isolated bound Tpm is given by the probability that a domain starting in state *1* does not end up in state *0*. We solved for the analytical solution of this probability by summing over all possible paths that start from state *1* and end in *0* then took the compliment. This resulted in the convergent solution:

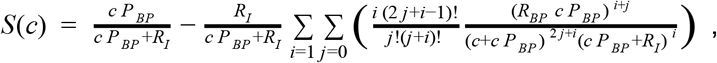

where *c* is the tropomyosin concentration, *B*_*BP*_ and *R*_*BP*_ are the sums of binding and release rate constants, respectively, for the edges of the tropomyosin domain (measured experimentally in Figure 2, See Table 1) and *R*_*I*_ is the rate constant for release of an isolated tropomyosin molecule from the actin filament (measured experimentally in Figure 4, See Table 1).

This model can be simplified by replacing the binding and release from the end of domains by the total elongation rate of *c P*_*BP*_ - *R*_*BP*_. This simplified scheme is shown in Figure 5A. The probabilities for each transition are shown below each reaction in Figure 5A. The probability of success is given by the product of the probability that a domain starting in state *1* moves all the way to the right of the chain. As all transitions after moving from state *1* to state *2* are equal to one, this probability is given by:

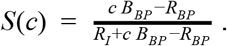

Comparing the probability of domain appearance calculated as a function of Tpm concentration using the full model and the simplified model shows that this simplification is valid for the process studied here (Supplementary Figure 9B).

### Implications of a fast release rate of isolated Tpm molecules from the actin filament

The value for the isolated Tpm release rate, *R*_*I*_, determined here from single-molecule data is possibly an underestimate (see Supplementary Figure 7). In this section, we discuss the implications of a faster release rate and present an alternative method to derive the ratio between the rate constants for binding and release of an isolated Tpm molecule on the basis of the experimental domain appearance rates.

In the limit where the isolated Tpm release rate, *R*_*I*_, is much faster than the total elongation rate (*c P*_*BP*_ - *R*_*BP*_), we can make the approximation that:

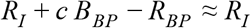

This approximation allows us to simplify the denominator for the probability of domain appearance, *S*(*c*), so that it becomes the following linear equation:

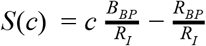

The domain appearance rate (given by *k*_*DA*_ = *c*.*B*_*I*_ .*S*(*c*)) divided by the concentration can then expressed as the following linear relationship:

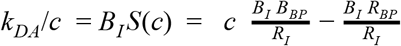

Supplementary Figure 10A shows the plot of the experimental data of *k*_*DA*_/*c* as a function of Tpm concentration. The experimental data is approximated reasonably by a straight line suggesting we are close to a regime where the release rate of an isolated Tpm molecule is dominant. Within this regime, the gradient of the best fit line (12.34 *nM* ^−2^*m*^−1^*s*^−1^) divided by the rate constant for binding at both edges of the Tpm domain (*B*_*BP*_ = 0.056 *nM* ^−1^*s*^−1^) gives the ratio of the ratio between the rate constants for binding and release of an isolated Tpm molecule (*B*_*I*_ /*R*_*I*_ = 220 *nM* ^−1^*m* ^−1^). It is notable that since the nucleation rate per nM is approximately linear the actual nucleation rate is approximately a parabola with an x-intercept at the critical concentration (Supplementary Figure 10B).

To explore how much faster the single Tpm dissociation rate, *R*_*I*_, needs to be in comparison to the total elongation rate (*c P*_*BP*_ - *R*_*BP*_), for the linear approximation to hold, we solved for the isolated Tpm binding rate (*B*_*I*_) using least squares as a function of the isolated Tpm release rate (*R*_*I*_). This curve is shown in Supplementary Figure 10C. For *R*_*I*_ above 10 *s*^−1^ there is less than a 4% percent discrepancy between the two methods. Above this value for *R*_*I*_, we expect the single Tpm binding to dissociation ratio (*n*/*r* = 220 *nM* ^−1^*m*^−1^) to be accurate.

### Supplementary Figures and Tables

**Supplementary Figure 1.**
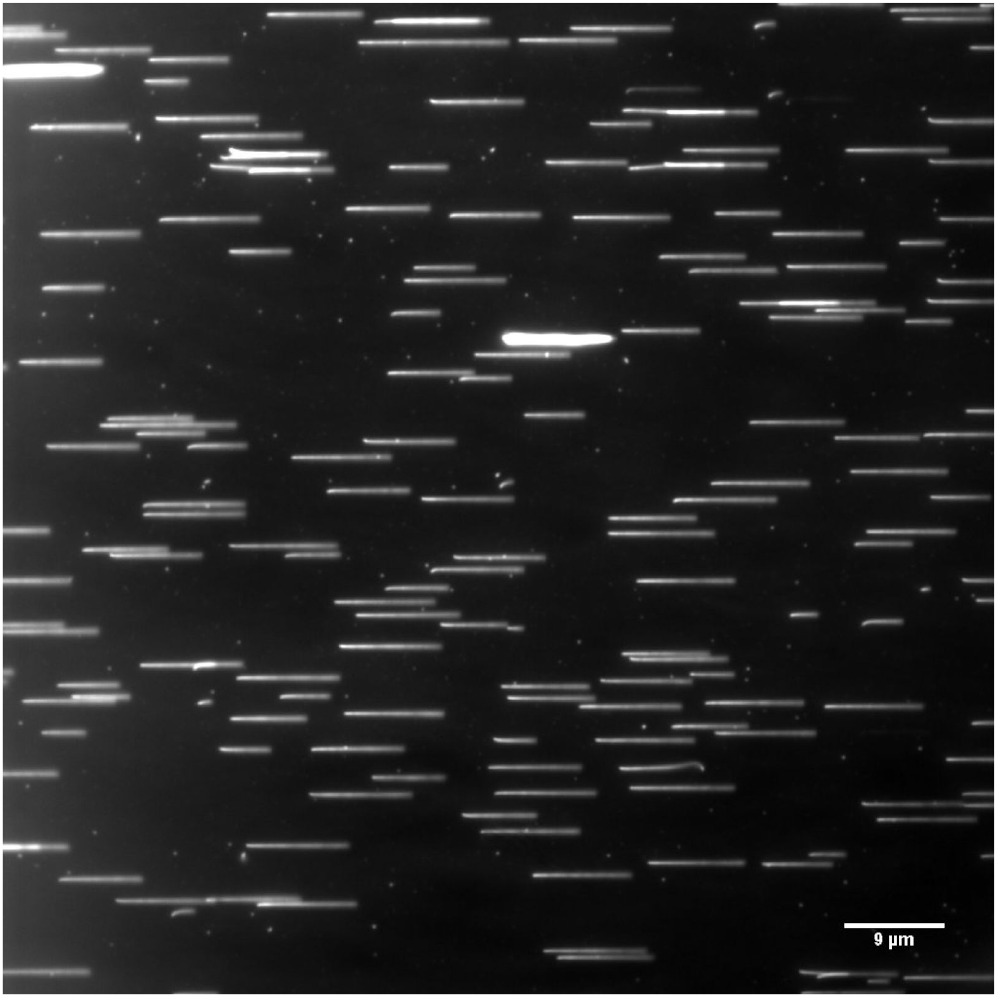
Density of actin filaments on surfaces. TIRF image showing actin filaments completely decorated by mNeonGreen-Tpm1.8 grown from surface-immobilised spectrin-actin seeds and aligned by the flow of buffer in the microfluidic channel. A field of view contained up to ~100 filaments for analysis.

**Supplementary Figure 2.**
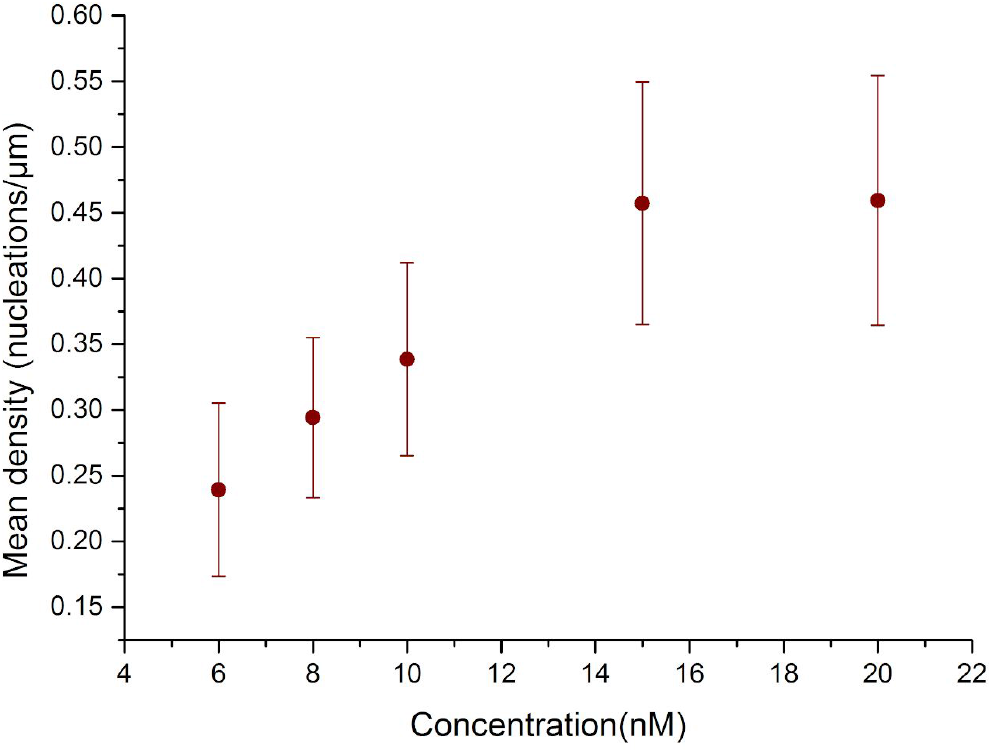
Dependence of the linear density of Tpm1.8 domain appearance on Tpm1.8 concentration. The number of Tpm1.8 domains that nucleate per unit length of actin filament increases with Tpm1.8 concentration at concentrations ≤15 nM. Points represent the mean and error bars represent the standard deviation; *N* (number of filaments) = 34 (6 nM), 31 (8 nM), 44 (10 nM), 43 (15 nM), 38 (20 nM).

**Supplementary Figure 3.**
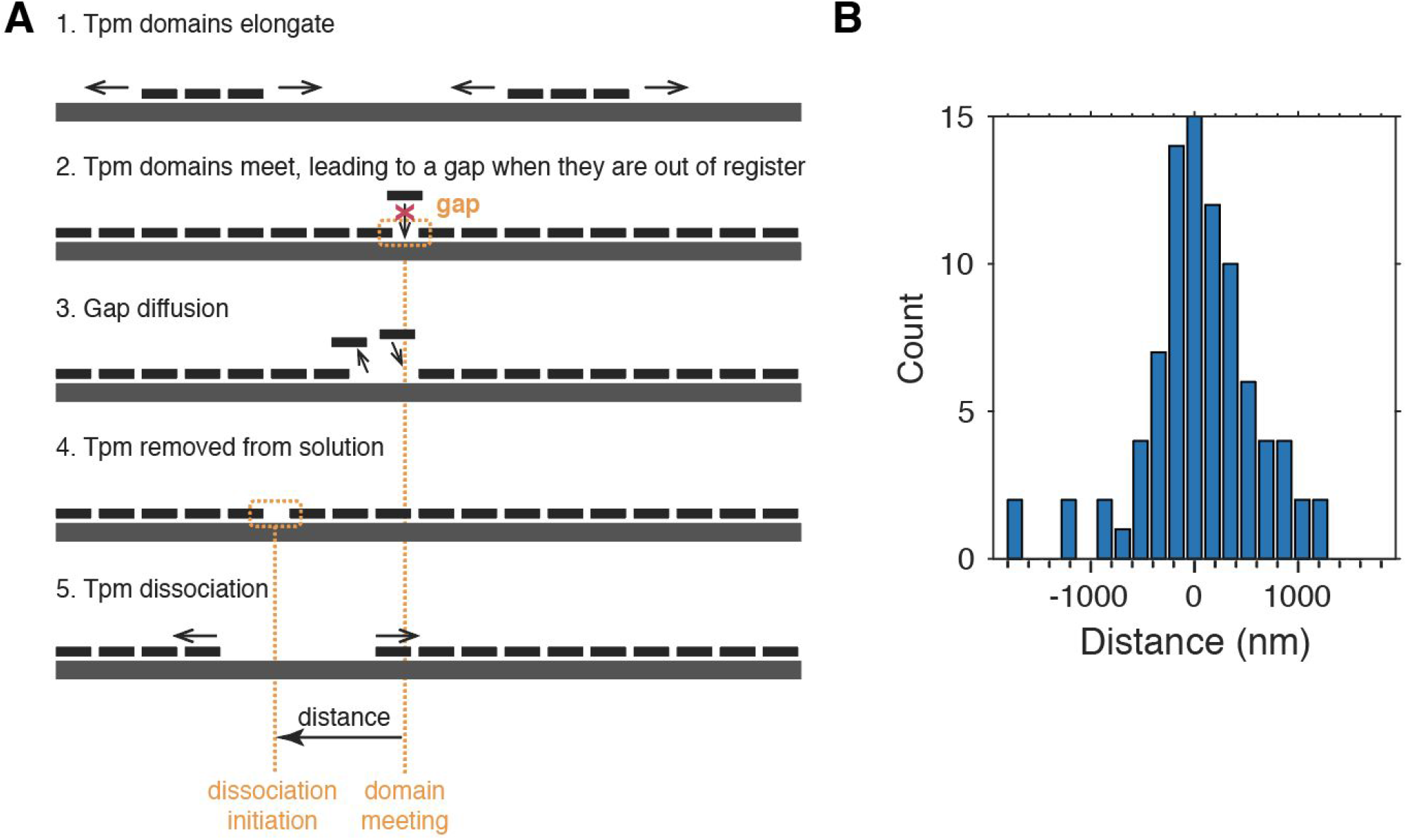
Relationship between points along the actin filament where two Tpm1.8 domains meet and points where dissociation initiates after Tpm1.8 wash-out. **A.** Schematic representation of the proposed molecular processes leading to the presence of gaps between Tpm1.8 domains during actin filament decoration that could serve as initiation points for Tpm1.8 dissociation after removal of Tpm1.8 from the flow channel. (1) In the presence of Tpm1.8 in solution, Tpm1.8 domains that have nucleated at different locations on the same strand of the actin filament elongate towards each other. (2) Each Tpm1.8 molecule covers 6 actin subunits such that two domains that have nucleated independently of each other at random locations of the actin strand can either be in register (in 1 out of 6 cases) or out of register (in 5 out of six cases) with each other. In the former case, the two domains can anneal into a single domain, whereas in the latter case a gap remains that is too short (between 1-5 actin subunits, i.e. 5.5-27.5 nm) to accommodate a Tpm1.8 molecule. (3) This gap between domains cannot be annealed and may diffuse in either direction along the actin filament as a result of Tpm1.8 dynamics at the domain edges. (4/5) When Tpm1.8 is removed from solution, these defects are likely to constitute points of instability leading to Tpm1.8 dissociation from the edges of the domains at both sides of the gap, which consequently shrink away from each other. **B.** Histogram of the distances between domain meeting points and the closest dissociation initiation points detected on the same filament; negative/positive values represent movement of the gap towards the pointed/barbed end, respectively. The distribution has a mean of −24 nm (i.e. it is approximately centered on zero) and a root mean square value of 557 nm, consistent with a model whereby dissociation initiates at gaps that have undergone diffusion towards either end of the actin filament. Methods: Domain meeting points were detected as those locations where the fluorescence signals of two domains elongating towards each other merge into a continuous line. We note that a gap smaller than the diffraction limit (~250 nm) would not appear as a discontinuity in the fluorescence signal, i.e. not detectable in this experiment. Dissociation initiation points were identified as points where gaps appeared in the fluorescence signal of the previously fully decorated actin filament. The histogram contains data from 48 filaments (18/30 filaments decorated in the presence of 5/10 nM mNeonGreen-Tpm1.8, respectively, before wash-out with buffer). We detected 174 domain meeting points during the binding phase of the experiments and 92 dissociation initiation points (53%). We then measured the distance between each dissociation initiation point and the closest domain meeting point on the same filament (5 dissociation initiation points could not be matched to a closest meeting point and are not included in the histogram). Note that gaps that diffuse all the way to either end of the actin filament disappear (i.e. “fall off” the end of the filament).

**Supplementary Figure 4.**
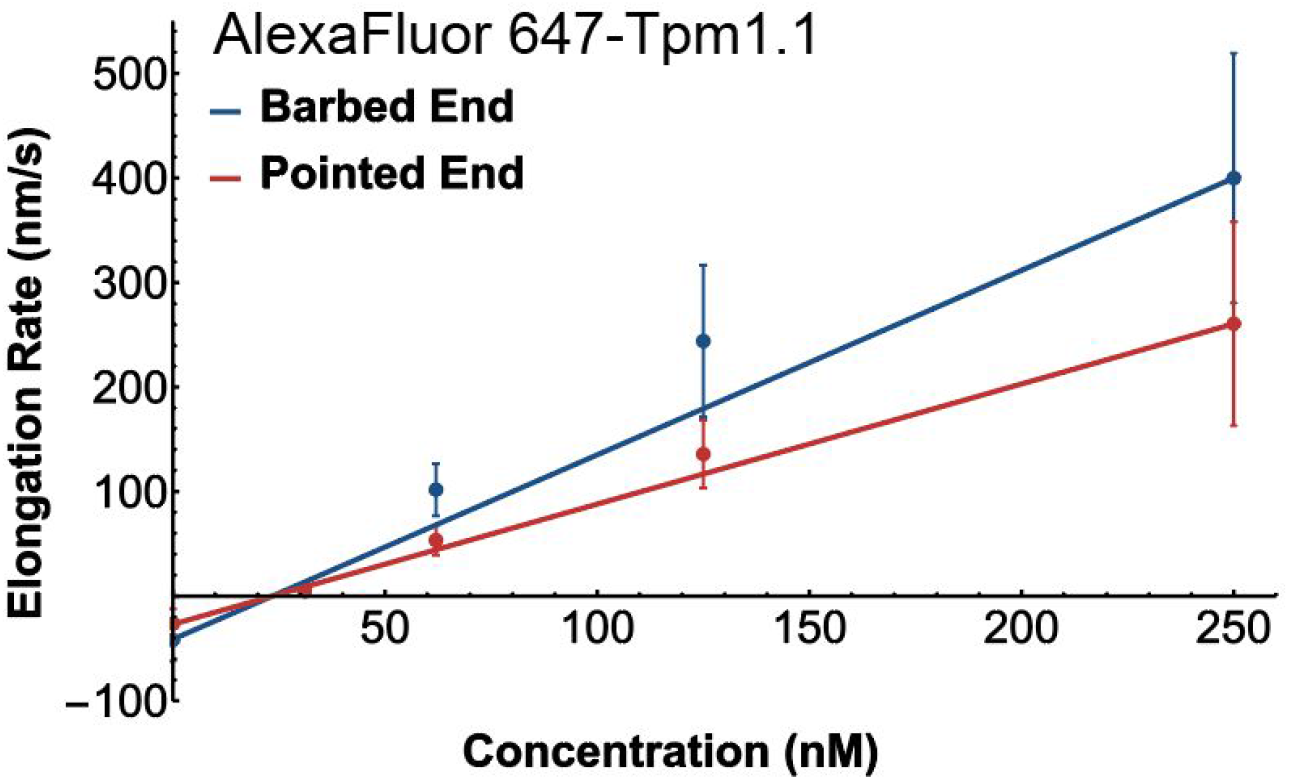
Asymmetric domain kinetics are observed with different Tpm1.1 labelling strategies. Concentration dependence of AlexaFluor 647-labelled Tpm1.1 domain elongation rates toward the barbed and pointed ends of the actin filament. The solid lines represent linear fits of the data, whereby the slopes give the elongation rate constants: 1.76 m.M-1.s^−1^ towards the barbed end and 1.14 m.M-1.s^−1^ towards the pointed end, giving a ratio of 1.54. For comparison, recombinant Tpm1.1 fused at its N-terminus to an alanine-serine extension and to mNeonGreen (mNeonGreen-Tpm1.1) showed the following elongation rate constants: 1.11 m.M-1.s^−1^ towards the barbed end and 0.667 m.M-1.s^−1^ towards the pointed end, giving a ratio of 1.66 (manuscript in preparation). This comparison suggests that fusion of a fluorescent protein to the N-terminus or chemical labelling of an internal cysteine results in labelled Tpm1.1 with similar assembly kinetics on actin, whereby the asymmetry in elongation and dissociation kinetics towards the opposing ends of the actin filament are maintained.

#### Methods

Recombinant Tpm1.1 with N-terminal alanine-serine extension was labeled by reaction with a maleimide derivative of AlexaFluor 647 as described previously (Nicovich et al (2016) Cytoskeleton 73(12), 729-738). Tpm1.1 contains a single cysteine residue at position 190, which is commonly used for labelling this isoform using thiol-reactive probes. Labelling conditions were chosen to achieve a labelling ratio of ~20% (i.e. 0.2 fluorophores per cysteine residue). Assuming random incorporation of the dye, this labeling ratio corresponds to a mixture containing 36% labeled dimers, whereby 89% of the labeled dimers contain one fluorophore and 11% contain two fluorophores. We have observed that higher labelling ratios perturb Tpm1.1 kinetics on actin suggesting that incorporation of two fluorophores per Tpm1.1 dimer affects binding to actin filaments. The tropomyosin isoforms Tpm1.1 and Tpm1.8 arise due to differential splicing of the transcript of the same gene, whereby Tpm1.1 contains exons 1a, 2b, 3, 4, 5, 6b, 7, 8 and 9a while Tpm1.8 contains exons 1b, 3, 4, 5, 6b, 7, 8 and 9d.

**Supplementary Figure 5.**
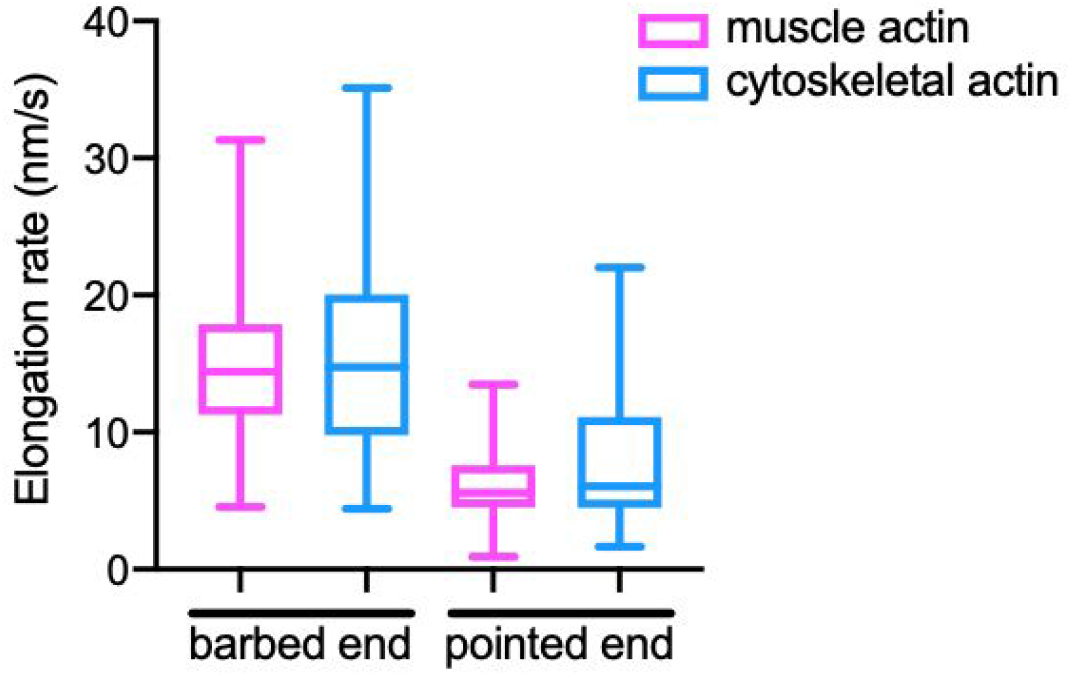
Tpm1.8 domain elongation rates towards the barbed and pointed ends of the action filament are similar on muscle and cytoskeletal actin. Actin filaments were grown from actin-spectrin seeds immobilised on the coverslip surface using either muscle actin or cytoskeletal actin. Subsequently, a solution containing 7.8 nM mNeonGreen-Tpm1.8 was injected into the flow cell and the decoration of actin filaments was imaged by time-lapse TIRF microscopy. Elongation rates towards the two ends of the actin filaments were determined from the slopes of Tpm1.8 domain edges in the kymographs obtained from the TIRF movies. Box plots showing the median and 25th and 75th percentile with whiskers extending from the minimum to the maximum data points; *N* = 81 slopes (barbed end, muscle actin), 34 slopes (barbed end, cytoskeletal actin), 72 slopes (pointed end, muscle actin), 24 slopes (pointed end, cytoskeletal actin).

**Supplementary Figure 6.**
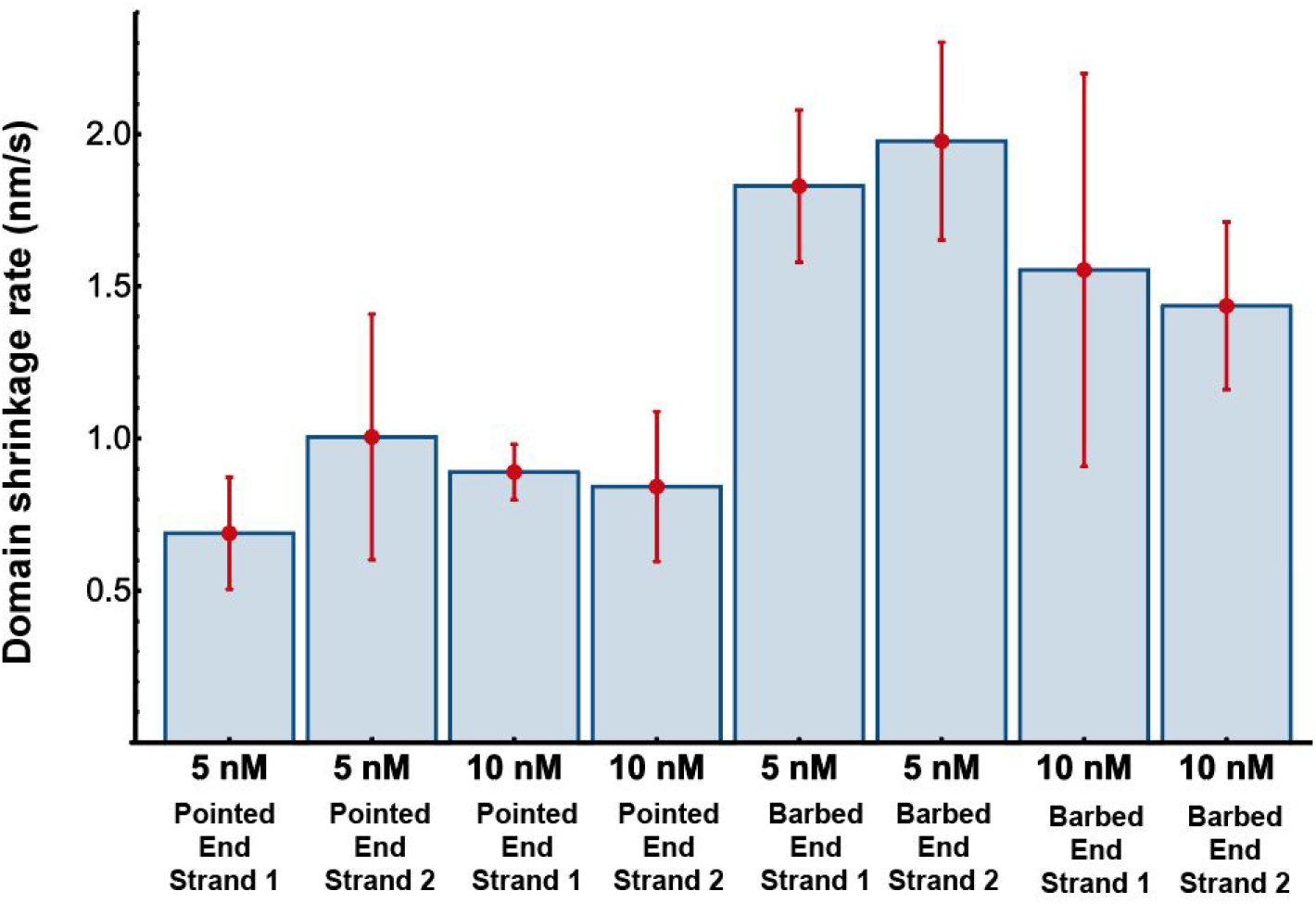
Tpm1.8 domain shrinkage kinetics. Comparison of domain shrinkage rate constants measured from the slopes in kymographs determined from actin filaments (decorated at either 5 nM or 10 nM mNeonGreen-Tpm1.8) after wash-out of free mNeonGreen-Tpm1.8 from the flow channel. Points represent the mean and error bars represent the standard deviation. Differences are not significant, except for pointed end versus barbed end values. Number of filaments: 42 (5 nM), 44 (10 nM); number of slopes: Pointed end, strand 1: 12 (5 nM), 4 (10 nM); Pointed end, strand 2: 25 (5 nM), 20 (10 nM); Barbed end, strand 1: 14 (5 nM), 6 (10 nM); Barbed end, strand 2: 42 (5 nM), 40 (10 nM). Comparison of pointed end rates: *p* = 0.76 (5 nM versus 10 nM) and *p* = 0.19 (strand 1 versus strand 2) using two way ANOVA (Tukey test). Comparison of barbed end rates: *p* = 0.14 (5 nM versus 10 nM) and *p* = 0.35 (strand 1 versus strand 2) using two way ANOVA (Tukey test).

**Supplementary Figure 7.**
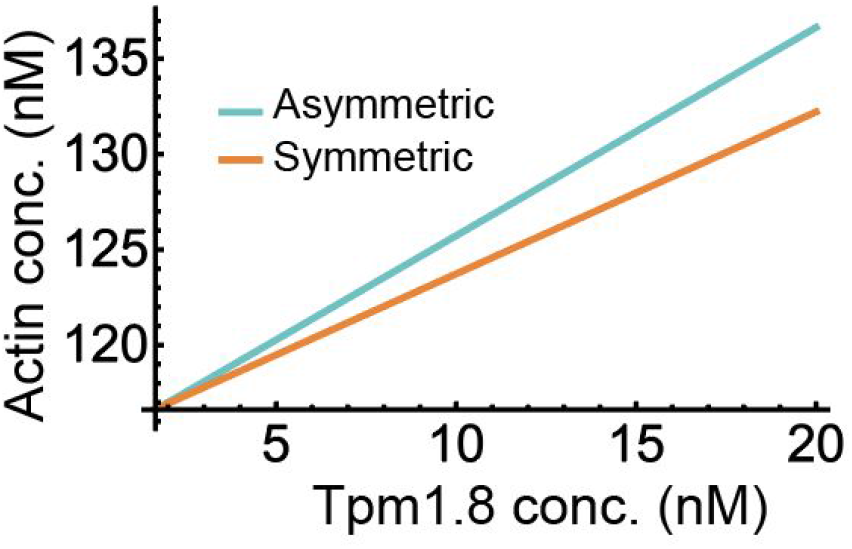
Plot of the maximum free concentration of actin for which Tpm1.8 binding outpaces actin polymerisation as a function of Tpm1.8 concentration for the experimentally observed Tpm1.8 elongation kinetics with asymmetry (green line) or without asymmetry (orange line, assuming that the rate constants for both edges are equal to the mean of the asymmetric rate constants). Actin polymerisation rates as a function of free action were calculated using published rate constants.

**Supplementary Figure 8.**
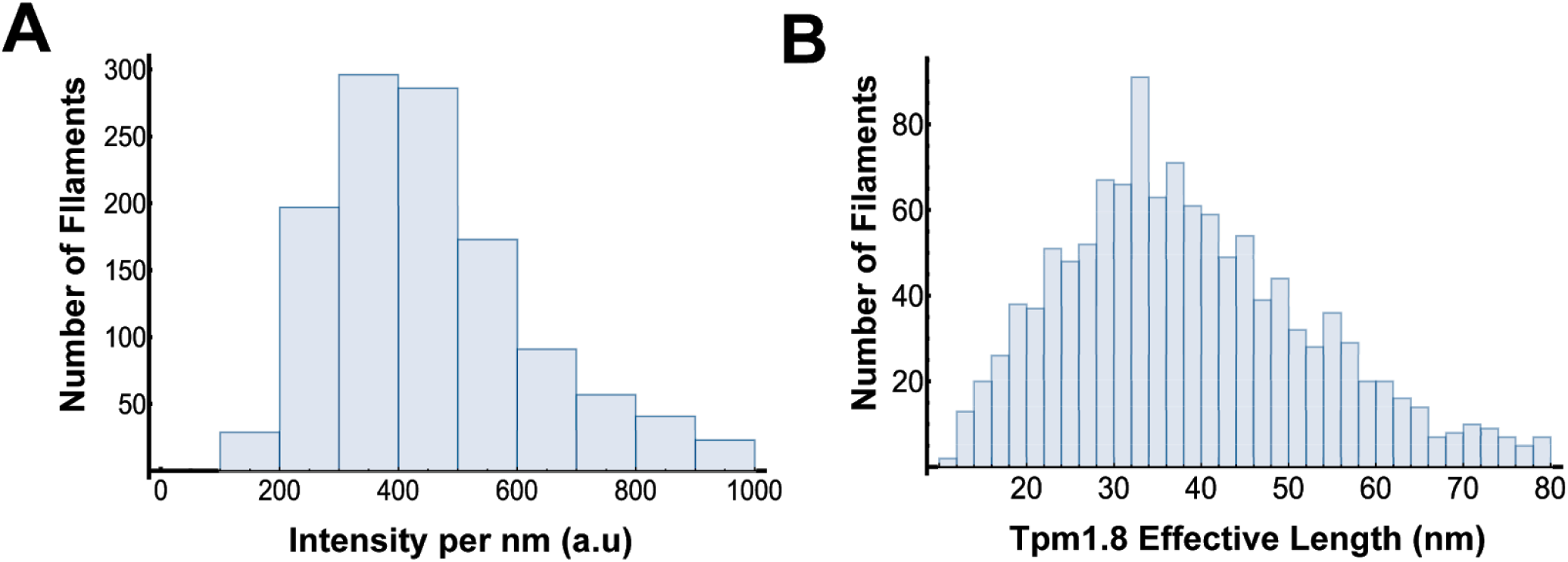
Calculation of the effective length of a Tpm1.8 molecule within a domain bound to an actin filament. **(A)** Histogram of the mNeonGreen-Tpm1.8 fluorescence intensity per unit length of a stretch of actin filament decorated (on one strand) with a continuous domain of mNeonGreen-Tpm1.8. **(B)** Distribution of distance covered by a single Tpm1.8 molecule (i.e. length added to a Tpm1.8 domain by addition of a Tpm1.8 molecule) with a mode value of 33 nm. The intensity of a single (dimeric) mNeonGreen-Tpm1.8 was determined by single-molecule photobleaching (Figure 4E).

**Supplementary Figure 9.**
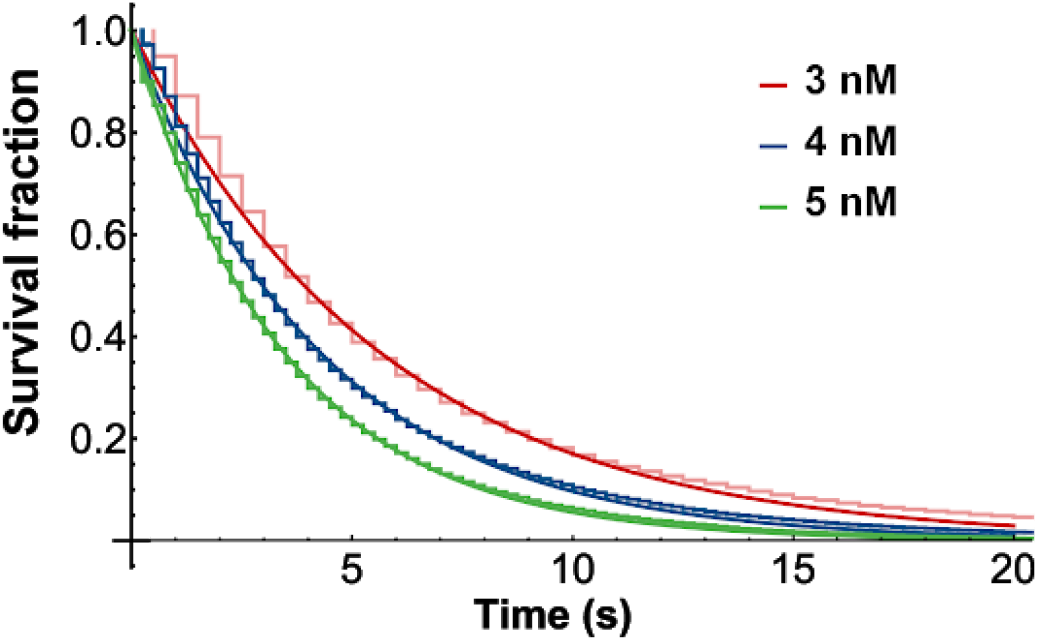
Calculation of the combined binding rate at both ends of a tropomyosin domain (*B*_*BP*_) from single-molecule data. Distributions of dwell times between positive steps in Tpm1.8 intensity traces measured at diffraction-limited domains in the presence of different Tpm1.8 concentrations. Curves were fitted with a single exponential function to obtain the rates (*c* × *P*_*BP*_) of Tpm1.8 binding to both ends of the growing domain for each Tpm1.8 concentration. The rate constant for binding at both ends of a tropomyosin domain (*P*_*BP*_, shown the Figure 4D) was calculated by dividing the rate obtained from the exponential fit by the respective Tpm1.8 concentration.

**Supplementary Figure 10.**
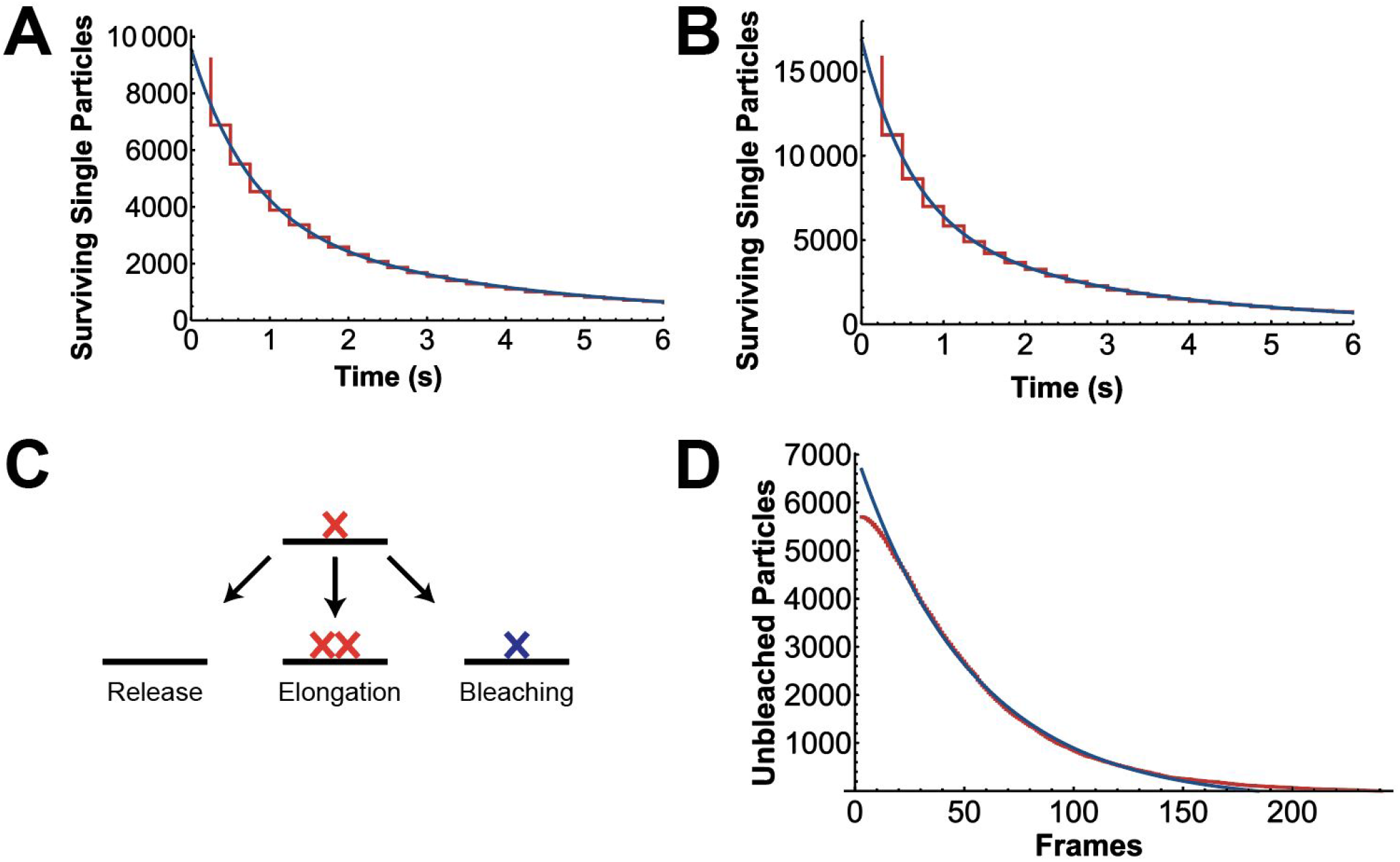
Estimation of the rate constant for release of an isolated tropomyosin molecule from the actin filament. **(A, B)** Survival functions of single-molecule lifetimes (red line) measured in the presence of 4 nM **(A)** or 5 nM **(B)** mNeonGreen Tpm1.8 in solution. The distributions were fitted with a double exponential decay (blue line). The fast process (rate constant of 1.34 s^−1^ [4 nM] and 1.65 s^−1^ [5 nM]) was attributed to the disappearance of single molecules, whereby the rate constant for release of an isolated tropomyosin molecule was obtained after subtracting the known rate constants for competing processes shown in panel C, i.e. photobleaching (obtained from the data in panel D) or addition of a second tropomyosin molecule (given by *c* × *P*_*BP*_, where *c* is the tropomyosin concentration and *P*_*BP*_ is the rate constant for binding at both ends of a tropomyosin domain). This analysis yielded an estimate for the rate constant for release of an isolated tropomyosin molecule of between 1–1.3 s^−1^. The slow process was consistent with the disappearance of signals from tropomyosin molecules that were misidentified as single molecules but likely part of small (partially bleached) tropomyosin domains. We note that the rate constant of the fast process is at the upper limit of rates that can be detected and measured with the frame rate used in the experiment (frame rate of 4 s^−1^ to avoid excessive photobleaching). As such, the estimated value represents a lower limit for the rate constant for release of an isolated tropomyosin molecule. **(C)** Schematic of the processes that lead to the disappearance of a spot with single-molecule intensity. **(D)** Survival curve of fluorophore lifetimes in single-molecule photobleaching experiments (red line). The rate constant (0.072 s^−1^) for photobleaching was obtained from a single exponential fit (blue line) of the survival curve.

**Supplementary Figure 11.**
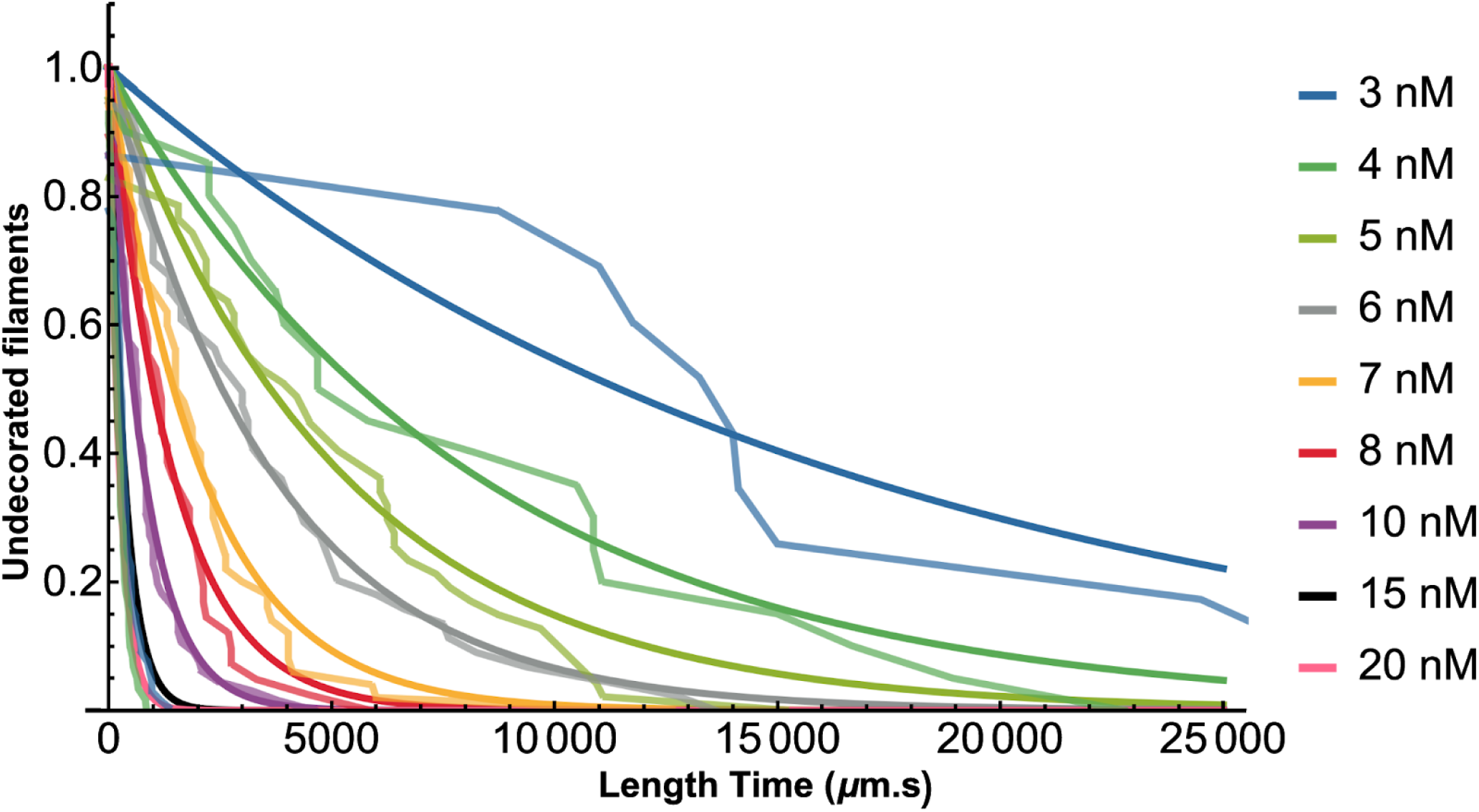
Calculation of domain appearance rates. Survival curves obtained by plotting the fraction of naked actin filaments (i.e. prior to the appearance of the first Tpm1.8 domain) as a function of the length×time constant. Fitting the exponentials gives the domain appearance rates. *N* (number of filaments) = 10 (3 nM), 19 (4 nM), 39 (5 nM), 42 (6 nM), 48 (7 nM), 38 (8 nM), 57 (10 nM), 52 (15 nM), 55 (20 nM).

**Supplementary Figure 12.**
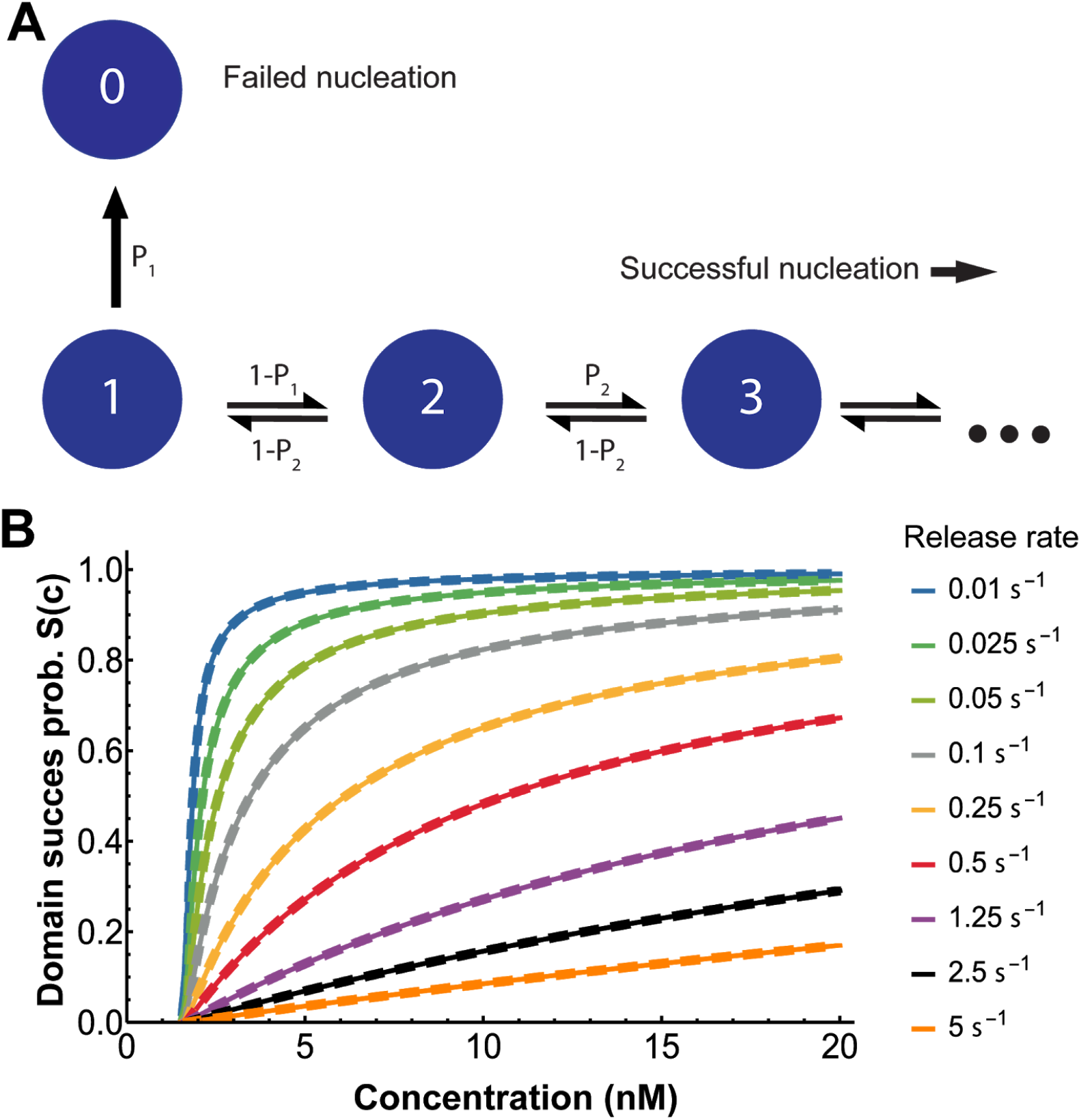
**(A)** A Markov chain depicting the probability of successful nucleation for the full model for domain appearance. Each state denotes the number of molecules in the domain. Isolated Tpm molecules bound to actin are in state *1*. Isolated molecules that dissociate move to the *0* state with probability *P*_1_. Alternatively, there is a probability, 1−*P*_1_, that a second molecule will bind to the isolated Tpm moving it to state *2*. For states containing more than one molecule, there is a probability *P*_2_ that it will become larger and a probability 1**−***P*_2_ that it will become smaller. These two probabilities are related to the reaction rates in Table 1 by *P*_*1*_ = *R*_*I*_/(*R*_*I*_+*c B*_*BP*_) and *P*_*2*_ = *c B*_*BP*_/(*R*_*BP*_+*c B*_*BP*_). **(B)** The domain formation success probability (*S(c)*) as a function of concentration for a range of different single tropomyosin release rates, *R*_*I*_. All curves eventually converge to 1 in the high concentration limit. Thus, at a sufficiently high concentration isolated bindings will essentially always develop into stable domains. However, the faster the monomer release rate, the higher the concentration needed to get equivalent levels of successful nucleation. The nucleation success probability calculated using the full model in (A) shown as continuous lines for each colour and the simplified model from Figure 5A (shown as dashed lines) overlay, showing the simplified model is valid within this range of values for *R*_*I*_.

**Supplementary Figure 13.**
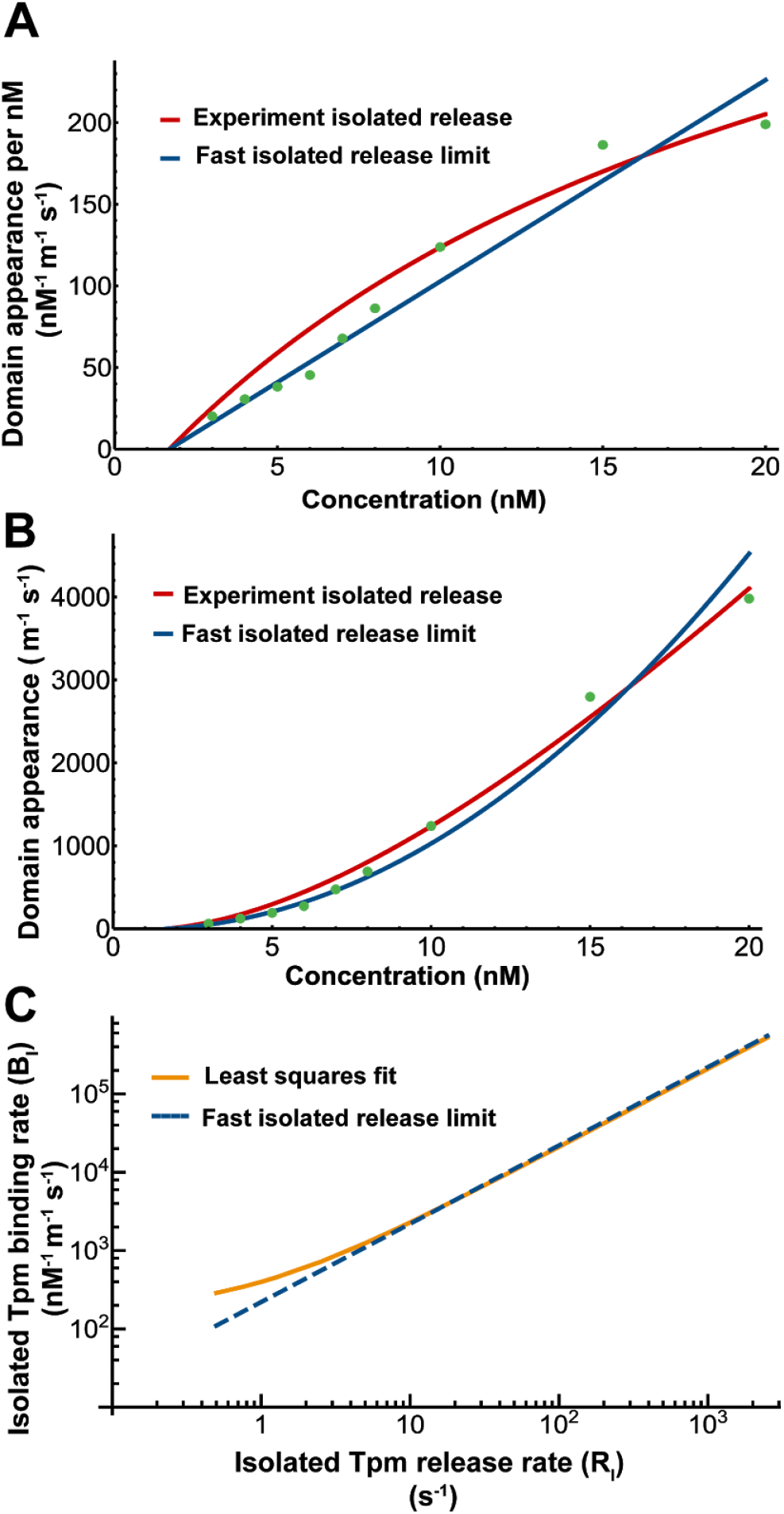
Nucleation with a fast isolated Tpm release rate. **(A)** Plot of the experimental domain appearance rate from Figure 5C divided by Tpm concentration (*k*_*DA*_/*c*) as a function of Tpm concentration (green dots). The best linear fit (constrained to pass through the x-axis at the critical concentration for Tpm1.8 assembly) is shown in blue (*k*_*DA*_/*c* = 12.34 nM^−2^ m^−1^ s^−1^ . *c* − 20.7 nM^−1^ m^−1^ s^−1^). The best fit curve using the experimentally measured release rate for an isolated Tpm molecule (*R*_*I*_ = 1.25 *s*−1) is shown in red. **(B)** Plot of the experimental domain appearance rate (*k*_*DA*_) as a function of Tpm concentration (green dots, from Fig 5C). Plot of the theoretical domain appearance rate as a function of concentration calculated by multiplying the fast release limit fit in (A) by the concentration at each point (blue), which yields a parabola with an x-intercept at the critical concentration. For comparison, the nucleation rate resulting from the best fit curve using the experimentally measured release rate for an isolated Tpm molecule (*R*_*I*_ = 1.25 *s*−1) is shown in red. **(C)** The best fit values for the binding rate constant for an isolated Tpm molecule (*B*_*I*_) as a function of the experimentally measured release rate for an isolated Tpm molecule (*R*_*I*_) shown in Red. If the single tropomyosin release rate is sufficiently fast (> 10 *s*−1), this relationship is approximately linear (shown by the dashed blue line) with a slope of 220 *nM* ^−1^*m* ^−1^ . This value is the ratio between the rate constants for binding and release of an isolated Tpm molecule.

**Supplementary Figure 14.**
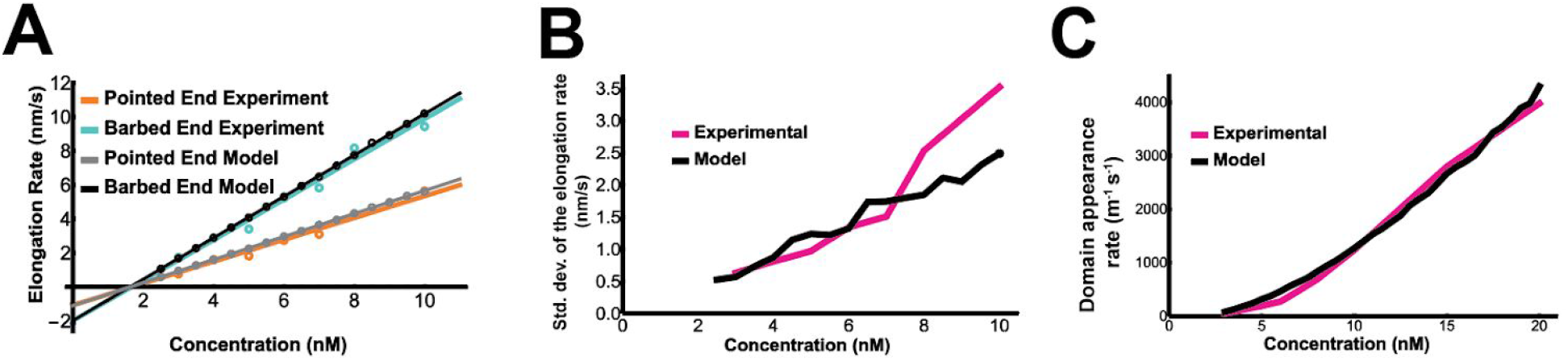
Comparison of parameters extracted from simulated and experimental kymographs of Tpm1.8 assembly on actin filaments. The model for Tpm1.8 assembly (see Figure 6A for interactions included in the model) was used to randomly generate 10^4^ kymographs, which were analysed using the same method as for experimental kymographs to extract kinetic parameters. **(A)** Elongation rates for Tpm1.8 binding towards the two ends of actin filaments obtained from experimentally obtained (orange/green; data and linear fit lines) or simulated kymographs (grey/black). **(B)** Variability of elongation rates as a function of Tpm1.8 concentration determined from experimentally obtained (magenta) or simulated (black) kymographs, calculated by taking the standard deviation of the residual of the linear least square fit. At concentrations >7 nM the variability increases more rapidly in experimental kymographs than in theoretical kymographs, possibly because it becomes increasingly difficult to accurately measure elongation experimentally with increasing concentration. **(C)** Nucleation rates as a function of Tpm1.8 concentration determined from experimentally obtained (magenta) or simulated (black) kymographs. The theoretical curve reproduces the overall shape of the experimental curve.

**Supplementary Table 1.** Asymmetric elongation is observed with different Tpm1.8 labelling and filament anchoring strategies. The ratio between Tpm1.8 domain elongation rates towards the barbed end versus the pointed end was consistently between 1.6 and 2.2, with the exception of 100% mCherry labelling. However, the asymmetry in elongation rate ratio was recovered using a solution of 50% mCherry-labelled and 50% unlabelled Tpm1.8. The data obtained using this mixture also allowed us to estimate an elongation rate ratio of 1.9 for unlabelled Tpm1.8. Taken together, these data suggest that the asymmetry in elongation is not an artifact due to the fluorophore or the anchoring strategy.

| Construct | Conc (nM) | Anchor | Elong. rate, barbed end ($\times 10^{-9} \text{ m} \cdot \text{s}^{-1}$) | Elong. rate, pointed end ($\times 10^{-9} \text{ m} \cdot \text{s}^{-1}$) | Rate const*, barbed end ($\text{m} \cdot \text{M}^{-1} \cdot \text{s}^{-1}$) | Rate const*, pointed end ($\text{m} \cdot \text{M}^{-1} \cdot \text{s}^{-1}$) | Elong. rate ratio |
| --- | --- | --- | --- | --- | --- | --- | --- |
| mNeonGreen-Tpm1.8 | 5 | gelsolin | 5.2±0.9 | 3.2±0.9 | 1.6 | 1 | 1.6 |
| mNeonGreen-Tpm1.8 | 20 | spectrin | 26.7±6.4 | 12.2±3.1 | 1.5 | 0.7 | 2.2 |
| mCherry-Tpm1.8 | 20 | spectrin | 24.2±5.1 | 22.2±4.6 | 1.3 | 1.2 | 1.1 |
| 50% mCherry-Tpm1.8+ 50% unlabelled Tpm1.8 | 20 | spectrin | 32.3±6.1 | 20.6±6.7 | 1.8 | 1.1 | 1.6 |
| mRubyII-Tpm1.8 | 7.8 | spectrin | 12.5±6.25 | 6.5±3.15 | 2 | 1.1 | 1.9 |

\*) The rate constant was estimated to facilitate comparison between conditions assuming a critical concentration of 1.65 nM in all cases.

